# Population genomics analysis with RAD, reprised: Stacks 2

**DOI:** 10.1101/2021.11.02.466953

**Authors:** Angel G. Rivera-Colón, Julian Catchen

## Abstract

Restriction enzymes have been one of the primary tools in the population genetics toolkit for fifty years, being coupled with each new generation of technology to provide a more detailed view into the genetics of natural populations. Restriction site-Associated DNA protocols, which joined enzymes with short-read sequencing technology have democratized the field of population genomics, providing a means to assay the underlying alleles in scores of populations. More than ten years on, the technique has been widely applied across the tree of life and served as the basis for many different analysis techniques. Here, we provide a detailed protocol to conduct a RAD analysis from experimental design to de novo analysis – including parameter optimization – as well as reference-based analysis, all in Stacks version 2, which is designed to work with paired-end reads to assemble RAD loci up to 1000 nucleotides in length. The protocol focuses on major points of friction in the molecular approaches and downstream analysis, with special attention given to validating experimental analyses. Finally, the protocol provides several points of departure for further analysis.

## 1. Introduction

Type II restriction enzymes have been powering significant aspects of population genetics since their discovery in the early 1970s (Kelly and Smith 1970; Smith and Welcox 1970). Each new generation of genome technology has been combined with restriction enzymes (RE) over the successive 50 years resulting in a continuously improving view of the distribution of genetic variation in natural populations. From assaying single loci with Restriction Fragment Length Polymorphisms (Botstein et al. 1980), combining REs with electrophoresis in polyacrylamide gels, to anonymously assaying whole genome sets of loci with Amplified Fragment Length Polymorphisms (Vos et al. 1995), combining REs with PCR amplification, to labeling all genomic loci with Restriction site-Associated DNA (RAD, Miller et al. 2007), combining REs with DNA microarrays. The trend accelerated with the advent of second-generation sequencing technologies, such as Illumina short-read platforms, that could sequence millions of small DNA fragments at low cost. The family of Restriction site-Associated DNA sequencing (RADseq) protocols (Davey et al. 2011; Andrews et al. 2016) has democratized the field of population genetics by enabling large-scale studies of natural variation (Benestan et al. 2015; Frugone et al. 2019), demographics (Marandel et al. 2020), population structure (Carlen and Munshi-South 2021), genetic maps (Amores et al. 2011; Mérot et al. 2021), landscape genetics (Dudaniec et al. 2018; Bay et al. 2021), methylation (Schield et al. 2016; Trucchi et al. 2016), and phylogenetics (Eaton et al. 2016; Near et al. 2018). As RADseq has matured over the last decade, several underlying processes affecting the application of the protocol and the resulting data have become better understood, including issues involving DNA quality and allelic dropout, and with that knowledge, the protocol has become a reliable instrument in the biologist’s toolkit.

In this manuscript we will present a full protocol for the analytical components of a RADseq experiment. After broadly introducing the set of RAD protocols (Section 1.1), we will discuss several important points, with respect to the molecular protocols, that often separate successful from failed experiments (Section 1.2). We then introduce the Stacks version 2 software (Section 1.3) and we present a full analytical protocol consisting of several major parts: how to set up your experiment and choose your enzymes via RADinitio simulation (Section 2.3), then, once the molecular library is created and sequenced (n.b., beyond our scope), we walk through cleaning and demultiplexing the data (Section 2.4), assembling RAD loci without a reference genome (Section 2.5), including optimizing *de novo* assembly parameters, assembling with a reference genome (Section 2.6), as well as a hybrid of the two methods (Section 2.7). We also demonstrate how to apply a population genetics frame to the data via a population-level analysis (Section 2.8). At the end of each analysis section, we focus on how to assay the success or failure of the underlying molecular library and analytical results. Finally, we provide four examples of exporting data outside of Stacks to further pursue a population genetics analysis, including assessing SNP and haplotype-based population structure, genetic divergence, and linkage disequilibrium.

### 1.1. The RAD family of protocols

A number of RADseq protocols have been introduced over the past decade (see Davey, et al. (2011) for an introduction and Andrews, et al. (2016) for an update), but they can be generalized into two broad categories, both of which have been successfully applied in many experimental contexts. For single-digest protocols (sdRADseq), DNA is extracted from an individual and is digested with a single restriction enzyme; a barcoded adaptor is attached, the DNA molecules are sheared to a uniform size, and the sample is then multiplexed into a library of dozens to hundreds of samples. Finally, the multiplexed library is PCR amplified to increase the fraction of RE-associated DNA from a small to a large fraction of the overall DNA in the library. We highlight a variant of this protocol, BestRAD (Ali et al. 2016) as it improves yield significantly by physically isolating the RE-associated DNA from the larger pool of DNA prior to sequencing.

Double-digest restriction enzyme protocols (ddRADseq, Peterson et al. 2012) define the second class. This group of protocols uses two restriction enzymes to isolate DNA loci from the genome – typically a rare cutting enzyme combined with a frequent cutting second enzyme – replacing the shearing step from sdRADseq. ddRADseq has a simpler implementation at the bench, however it introduces some additional analytical challenges. The choice of enzyme combinations can be useful when working with very large genomes to select the right subset of DNA loci to work with, while imprecise size selection, in conjunction with dual enzymes, can result in loci being variably present across different individuals. In addition, PCR duplicates cannot be identified in a ddRADseq analysis unless molecules are labeled with a random oligo sequence as part of the adaptor (Hoffberg et al. 2016).

### 1.2. Top molecular impediments to good results

Bioinformatic data analysis is itself rarely fruitful unless it is preceded by the successful execution of a molecular protocol at the bench. Our experience in designing and implementing Stacks and subsequently in RADseq data analysis, has highlighted three aspects of molecular library construction that are critical to achieving a successful experimental analysis. First, a sufficient amount of DNA must be available from each sample to be included in the molecular library. In the hands of an experienced technician, and with a thorough understanding of the tradeoffs, a successful RAD library can be prepared using small amounts of DNA. More frequently, however, limited amounts of DNA will cascade negatively into later parts of the protocol, increasing the likelihood of library construction failure – for a fraction of the samples, or for the entire library. For this reason, RADseq experimental design should emphasize maximizing DNA input.

Second, and related to the DNA input, are the effects of PCR amplification on a library. In any sequencing library, the goal is to obtain many copies of the host DNA – by extracting it from many cells in the organism – this will provide a high amount of information with respect to which alleles are present to enable genotyping. Nearly all RAD protocols include an amplification step to increase the prevalence of RE-associated DNA above the remaining genomic DNA (which will not be sequenced). PCR is very effective at increasing copies of template DNA molecules; however, if too few templates are present from the host genome, then the PCR process will instead amplify copies of the limited templates. These PCR duplicates do not add information with respect to the underlying alleles in the host genome – if an allele was not initially sampled it will remain missing after PCR amplification – and they will definitely consume sequencing capacity. So, amplification can present the illusion of high information content. Sequencing a library rich in PCR duplicates yields a reduced amount of informative DNA sequence from which to genotype. Sequencing such a library to very high depths will slightly increase the amount of informative sequence but ultimately, the library cannot provide the information it never contained due to a lack of template DNA. And while it may be tempting to associate these affects with the number of PCR cycles used (and to therefore adjust cycles down), ultimately it will not solve the problem. Thus, the amount of DNA present should be assayed independently of amplification.

Third, and still the most common problem we see in the Stacks analysis help forum is a lack of sequencing coverage. Fortunately, this is a problem that is eminently preventable with the proper planning. But for a successful analysis, a RAD practitioner should aim for high coverage with the ability to accept lower coverage in case of molecular difficulties. Therefore, the RAD practitioner should not try to over-economize a library, which can end up producing coverage too low to reliably call polymorphisms and yielding a homozygous bias in those polymorphisms that are called. A practitioner ideally should plan for a minimum of 20-30x coverage per individual in the sequenced library.

### 1.3. Stacks version 2: *de novo* and reference-based RAD analysis

Stacks was one of the first analysis pipelines released to work specifically with RAD data – initially for the *de novo* creation of genetic map markers (Amores et al. 2011; Catchen et al. 2011), and then expanded to work with both *de novo* and reference-aligned data more generally for any population genetics analysis (Catchen et al. 2013). More recently, a major update (version 2) was made expanding the software to work specifically with paired-end reads, using the single- and paired-end reads to construct individual RAD loci up to 1 kilobasepair (kbp) in size (Rochette et al. 2019).

Stacks is architected to execute in three major phases: first data are demultiplexed, breaking the molecular library back down into individual samples, the existence of the RE cutsite remnant is verified, and the data are cleaned (low quality reads are discarded). Second, the main pipeline executes. If data are to be assembled *de novo* – without a reference genome – then the single-end reads are assembled into loci in each sample independently (`ustacks`). These loci are merged together, matching homologous loci across samples, in the construction of a metapopulation catalog (`cstacks`), and samples are then independently matched to that catalog (`sstacks`). Next, data are reorganized by the `tsv2bam` program so that loci from all samples are in the same order (loci start off in an arbitrary order among samples) to increase analytical efficiency. Finally, `gstacks` executes, assembling a paired-end contig in the metapopulation for each locus. It does this for each locus by taking the single-end reads used to assemble that locus and identifying the corresponding set of paired-end reads. These reads are collected across the metapopulation, down-sampled to a reasonable number (∼1,000 by default) and fed into a *de bruijn* graph assembly algorithm to form the paired-end contig. The paired-end contig is then merged with the single-end locus, using a suffix tree algorithm, and all reads from all samples in the metapopulation are then aligned to the resulting locus. Single Nucleotide Polymorphisms (SNPs) are then called, using a Bayesian model-based approach. First, the presence of a polymorphism is detected, using all reads in the metapopulation. This likelihood then serves as the Bayesian prior to genotype each sample individually at that locus. In the case of reference-aligned data, the user aligns their processed reads to a genome (with e.g. BWA, Li 2013) and the main pipeline starts directly at the `gstacks` program.

A third, hybrid mode to run the main Stacks pipeline exists: one can take loci assembled *de novo*, align them to an arbitrary genome after assembly, and then inject those genomic coordinates back into the dataset (we show this *integration* method in Section 2.7).

Finally, given the output of `gstacks`, the final component of the pipeline to run is the `populations` program, which will consider the resulting `gstacks` genotypes in a particular biological context. Given a *population map* (a list of sample names and the populations they belong to) the `populations` program will calculate summary statistics (e.g., π, F_IS_, loci that are in Hardy-Weinberg Equilibrium), divergence statistics (e.g., D_XY_, and F_ST_), as well as produce a number of exports for further analysis (e.g., VCF, STRUCTURE, and/or GenePop to name just a few). The `populations` program is designed to be executed multiple times on the same assembled and genotyped catalog, adjusting filters each time. Many filters are available, including a minor allele frequency filter, a population frequency filter, etc. The biological frame can also be modified (without changing the core genotypes calculated previously) by running `populations` with a different population map to calculate statistics in different ways (e.g., dividing the individuals into populations by geography, sex, or phenotype).

### 1.4. Stacks haplotypes versus SNPs

One of the strongest aspects of RAD-derived data is that a RAD locus can easily be phased. That is, the alleles from the underlying SNPs can be matched up into haplotypes. This was one of the drivers behind the construction of paired-end contigs with Stacks version 2, and it enables haplotypes of up to 1kbp to be reconstructed from each RAD locus in the metapopulation (locus length is dependent on the size selection step of the underlying molecular protocol). These haplotypes are useful for estimates of nucleotide diversity (π), for input into phylogenetics models, the estimation of divergence statistics, such as D_XY_, and F_ST_, and the inference of coalescent genealogies. The `populations` program produces haplotype-level statistics alongside the SNP-based statistics (e.g., Φ_ST_) and it also provide several haplotype-specific filters (e.g., `-- filter-haplotype-wise` will filter rare SNPs from haplotypes to decrease the amount of missing data).

## 2. Methods

### 2.1. Pre-sequencing: RADseq experimental design

Before sequencing, experimenters should seek to understand at least the base differences between protocols. Each protocol has its own molecular and experimental biases and can produce different magnitude and quality of data in different experimental systems. This means that, first, not all RADseq or RAD-like protocols are interchangeable and, second, that the success of an experiment using the technology is strongly dependent on the selection and implementation of a molecular protocol. Moreover, the problem is exacerbated by similar (and often uninformative) names of some of the protocols (Campbell et al. 2018), for example, “Genotype-by-sequencing” or GBS is both a specific protocol and a generic reference to RADseq-like experiments.

Before protocol selection, researchers should determine what genomic resources are available for their system of study, including the most closely related species with a reference genome (the NCBI Assembly organism browser is a good resource, https://www.ncbi.nlm.nih.gov/assembly/organism/). This reference can be used for the direct estimation of RADseq loci, in addition to being a resource for reference-based analyses. Researchers should also check the literature for any available genomic studies in the system (or in closely related organisms) to determine if RADseq has been previously applied to the system, verifying the specific protocol used and its implementation. The results of these previous publications can be used as guidelines from which to design a new experiment.

To begin planning, a researcher should focus on the tradeoffs between the number of samples they want to include in the experiments, the number of RAD loci (and variant markers) they want to produce from those samples, and the required sequencing capacity to achieve those goals. The number of loci generated varies from species to species and from protocol to protocol. The right number of markers to obtain depends on the biological questions being asked. For example, an assessment of population structure might be performed with a few thousand independent sites of the genome and might allow for the sequencing of fewer sites across more individuals. In contrast, a genome scan might need tens of thousands of makers evenly distributed along the genome and might require a marker-dense RAD protocol. Sequencing requirements are also tightly linked to the estimation of RAD loci. Choosing a protocol which produces a few markers in a library of a few individuals might result in over-sequencing the selected markers, while a very marker-dense library might result in under-sequencing if too many samples are added in the library and sequencing capacity is not scaled accordingly.

Below we show examples of how to calculate the expected number of RAD loci from a given reference genome, followed by an example of how to calculate sequencing capacity (in the form of number of samples and coverage) from these loci estimates.

### 2.2. Data set used in this protocol

In this protocol, we will be using a RADseq dataset from two populations of the Patagonian Blenny, *Eleginops maclovinus*. This fish species is native to the coastal regions of southern South America and is an outgroup to the cold-specialized radiation of Antarctic notothenioid fish (Chen et al. 2019). Sixty individuals from two populations in southern Chile (Valdivia n=37, Puerto Natales n=23) were processed into a single-digest RAD library (Baird et al. 2008; Etter et al. 2012) generated using the RE *SbfI*. The library was sequenced (2×150bp) on a single lane of an Illumina NovaSeq6000 at the Roy J. Carver Biotechnology Center sequencing facility at the University of Illinois, Urbana-Champaign.

A chromosome-level genome assembly was additionally generated for the species using long molecules from a PacBio Sequel 2 sequencer, which were then scaffolded using Hi-C. The assembly contains a total of 628 sequences, including 24 chromosome-level contigs and smaller unplaced scaffolds, for a total size of 613Mb and a scaffold and contig N50 of 26.7Mb and 7.6Mb, respectively.

All data analyzed in this protocol, as well as shell scripts to run the analysis and R scripts to generate summary statistics and plots, are available online at: https://bitbucket.org/CatchenLab/mimb-stacks2-protocol.

### 2.3. Simulations

In designing an experiment, we need to understand the architecture of the genome of our species of interest, how many RAD loci particular enzymes will yield, and how much sequencing will be required to assay those loci in the populations we wish to study. The first step is to determine the effects of RE and RAD protocol of choice.

#### 2.3.1. Calculating an estimated tally of RAD loci

To estimate the number of RAD loci we will tally the number of recognition sequences of one or more restriction enzymes in a reference genome. For sdRADseq, each enzyme recognition sequence will yield two tags, one from each DNA strand, while double-digest protocols require the recognition sequences for both enzymes to be within a certain insert-size window. The number and distribution of these RE recognition sequences are dependent on the size of the genome, its overall GC content, and the distribution of di- and tri-nucleotides in the sequence (Herrera et al. 2015). However, the number of RAD loci experimentally recovered can deviate from the number of observed cut sites. For example cut sites in close proximity may interfere with one another, reducing coverage (Davey et al. 2013).

There are several software packages designed around this initial estimation of RAD loci, including SIMRLLS (Eaton et al. 2016), SIMRAD (Lepais and Weir 2014), DDRADSEQTOOLS (Mora-Márquez et al. 2017), DDRAGE (Timm et al. 2018), PREDRAD (Herrera et al. 2015), and RADinitio (Rivera-Colón et al. 2021). All these programs base their estimation on counting restriction enzyme cut sites on a reference sequence but vary in the way they account for the variation introduced by the presence of repetitive loci, natural polymorphism, and technical error. For our estimates, we will be using the software RADinitio, as it can model the effects of genetic variation, protocol and enzyme choice, PCR duplicates, and sequencing coverage affecting the RADseq data generated.

Since there is a reference assembly available for *E. maclovinus*, we can perform the estimation of RAD loci directly. First, RADintio requires the reference genome in FASTA format, as well as a list of the sequences to simulate. The list of sequences can be obtained directly from the FASTA file by first extracting all the sequence ID lines and removing the ID identifier ‘> ‘ alongside any comments after the FASTA ID, saving them to a new file:

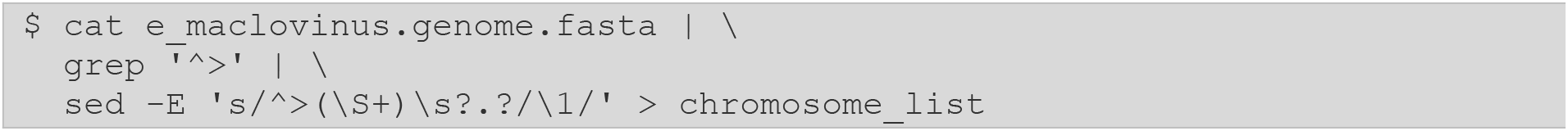

*Note! As we will be presenting shell commands in this protocol, we will symbolize the command line prompt with the ‘*`$` *’ character. This character is not part of the command but signifies the prompt at which the reader will type the command. To make our commands more readable in this limited space, we will divide them across lines using the backslash character, ‘*`\`*’. If the reader is typing commands all on one line, ‘*`\` *’ is not part of the command itself and should be omitted. Otherwise, ‘*`\` *’ tells the shell to continue the command onto the next line, instead of executing it*.

The file `chromosome_list` will now contain a list of all the chromosomes, one per line, which in the case of the *E. maclovinus* assembly are named as `HiC_scaffold_X`. We can look at the first 10 lines of the file using the `head` command.

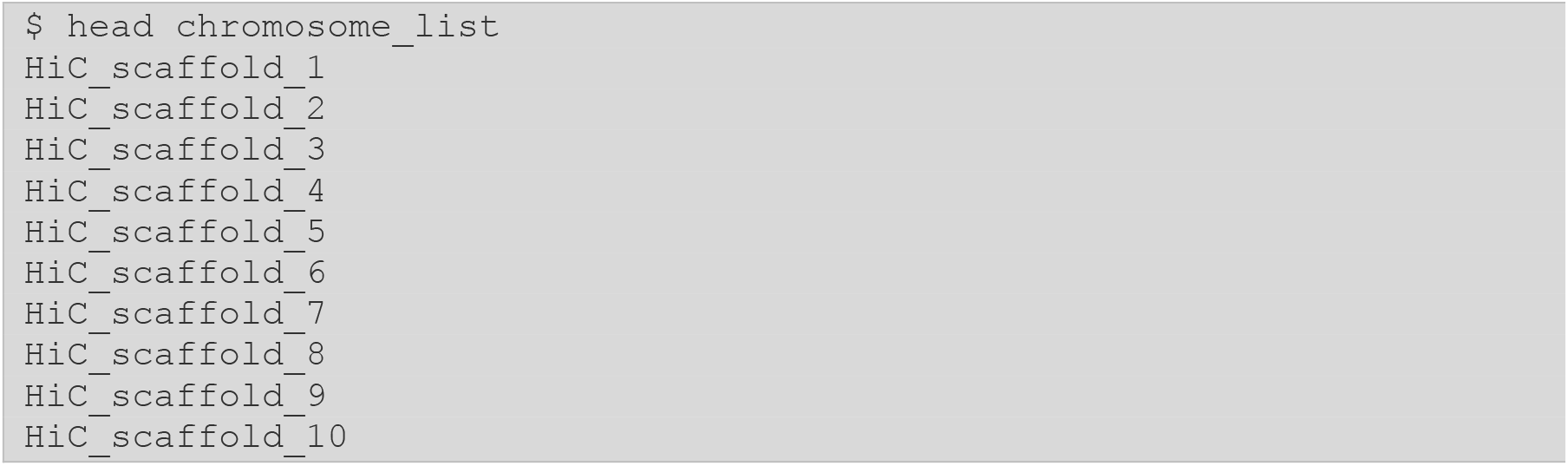

Since we want a tally of all possible RAD loci in the genome, we will pass all the available sequences to RADinitio. However, this chromosome list file can be edited if simulating over a subset of specific sequences is needed.

Now that we have the list of chromosomes, we can run `radinitio --tally-rad-loci`. The software requires the path to the reference genome in FASTA format (`--genome`), the path to the text file containing the list of chromosomes (`--chromosomes`), the path to an output directory to write the results (`--out-dir`), the type of library type to simulate (`--library-type`), and the RE (`--enz`). In this instance, we are simulating a single-digest RAD library using the enzyme *SbfI*. Additional options to simulate double-digest libraries, additional enzymes, and controlling insert sizes can be found in the RADinitio manual (http://catchenlab.life.illinois.edu/radinitio/manual/).

First, we will create a new output directory, which we will name according to the type of library being simulated. This will make it easier to identify the contents of the directory once we simulate libraries across several parameters.

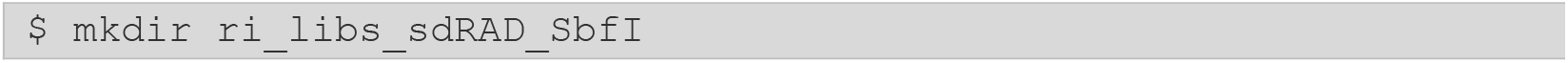

Next, we will run RADinitio using the parameters specified above. When reading this command remember that the ‘`\`’ character is a special escape character that allows us to continue the command on the next line, making longer commands easier to read. Additionally, the ‘`#`’ character is used to specify comments, meaning the text after it will not be interpreted as part of the command. These comments are a useful way to annotate your code, helping describe the purpose of specific options.

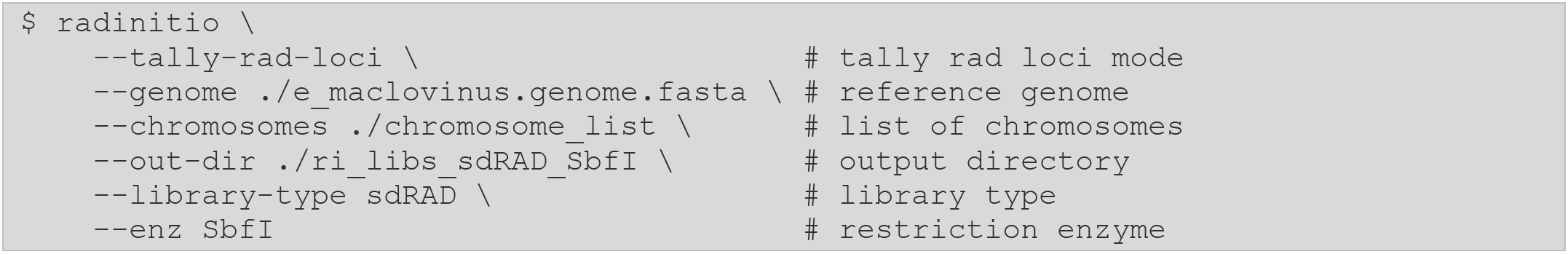

If we were to type this same command in a single line without using any comments it would look like this:

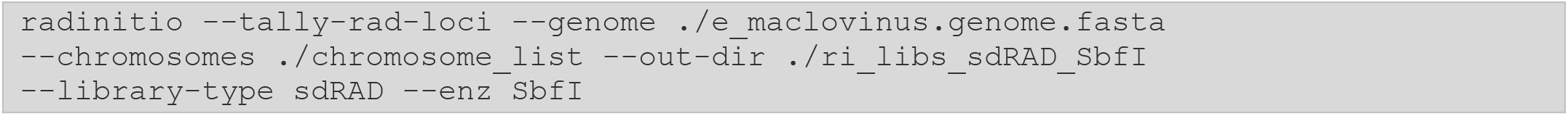

Executing the command simulates the *SbfI* single-digest library on the *E. maclovinus* reference genome. The produced tally of loci can be found in the `radnitio.log` file. We can observe that RADinitio found 52,644 RAD loci.

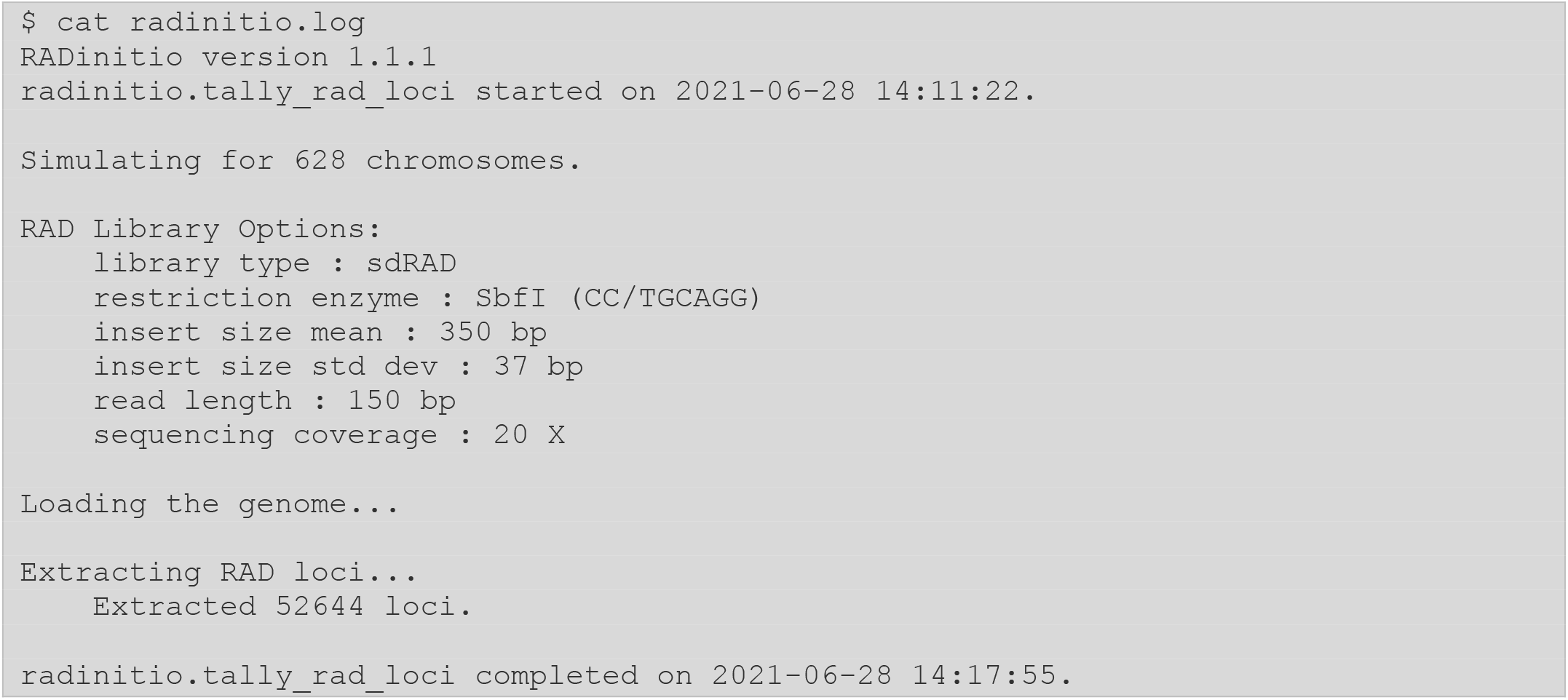

Note that the `--tally-rad-loci` command only accounts for variation in the proximity of loci and the effects of insert size when performing the tally of loci. More complex simulations, such as those introducing the effects of population polymorphism and sequencing can be performed with other RADinitio options (such as `--simulate-all).`

When no reference genome is available, a reference from a closely related species, for example one in the same genus or family, can substitute to estimate the number of RAD loci. These heterospecific genomes can also serve as an anchor for loci generated *de novo* using an *integrated* approach (Section 2.7). In the case that no single close reference is available, a good strategy is to perform simulation on several references to bound the expected results and ameliorate the effects of the evolutionary distance between the organism of study and the available references. Note that, when making these estimates, it is less important to have an exact count of the loci in the genome – after all any reference genome is bound to be different from the sequenced individuals – and more important to have an estimate of the magnitude of data expected.

To elaborate on the process described above, we performed several simulations and estimates of recovered RAD loci using various library protocols on seven different taxa (Table 1). We simulated both single- and double-digest loci using a variety of enzymes with varying GC content in their recognition sequences. For sdRADseq we used the GC-rich 8 bp cutter *SbfI* (CC/TGCAGG), the GC-rich 6 bp cutter *PstI* (C/TGCAG), and the AT-rich 6 bp cutter *EcoRI* (G/AATTC). For double-digest RAD, we used these three main cutters (*SbfI, PstI*, and *EcoRI*) in combination with the AT-rich 4 bp *MseI* (T/TAA), the GC-rich 4 bp *MspI* (C/CGG), and the 4 bp *AluI* (AG/CT) which contains 50% GC in its recognition sequence.

**Table 1:** Tally of RAD loci based on RADinitio simulations.

| Protocol | Enzyme | *Heliconius*<br>butterfly | *Mimulus*<br>Monkeyflower | Swainson's<br>Thrush | Threespine<br>Stickleback | Patagonian<br>Blenny | S. Georgia<br>Icefish | Tasmanian<br>Devil |
| --- | --- | --- | --- | --- | --- | --- | --- | --- |
| sdRAD | *SbfI* | 2,050 | 2,328 | 134,310 | 38,666 | 52,644 | 70,432 | 67,676 |
| sdRAD | *PstI* | 53,452 | 77,168 | 1,894,498 | 549,448 | 868,478 | 1,074,684 | 1,154,034 |
| sdRAD | *EcoRI* | 216,792 | 135,520 | 690,630 | 144,204 | 209,284 | 287,342 | 2,458,596 |
| ddRAD | *SbfI-MspI* | 318 | 337 | 17,844 | 10,059 | 7,939 | 14,853 | 10,403 |
| ddRAD | *SbfI-MseI* | 1,558 | 1,398 | 58,227 | 20,377 | 31,135 | 40,080 | 37,403 |
| ddRAD | *SbfI-AluI* | 753 | 916 | 78,856 | 17,949 | 25,495 | 32,548 | 32,858 |
| ddRAD | *PstI-MspI* | 6,474 | 10,745 | 171,384 | 135,973 | 123,178 | 208,906 | 116,396 |
| ddRAD | *PstI-MseI* | 42,833 | 47,505 | 960,386 | 297,370 | 519,804 | 629,062 | 696,153 |
| ddRAD | *PstI-AluI* | 18,184 | 32,178 | 1,103,549 | 254,298 | 419,391 | 493,069 | 540,275 |
| ddRAD | *EcoRI-MspI* | 20,589 | 17,746 | 49,055 | 34,058 | 27,914 | 54,182 | 82,368 |
| ddRAD | *EcoRI-MseI* | 179,728 | 90,052 | 388,664 | 80,593 | 129,999 | 171,451 | 1,681,957 |
| ddRAD | *EcoRI-AluI* | 73,855 | 49,573 | 373,678 | 65,770 | 100,383 | 129,502 | 1,029,578 |

We observe several important patterns. First, a given enzyme (or combination of enzymes) does not produce the same magnitude of loci across all genomes. Notice how *EcoRI* yields 2.5 million loci in the Tasmanian devil (*Sarcophilus harrisii*, genome size 3.08 Gbp), but only 136k in the bush monkeyflower (*Mimulus aurantiacus*, genome size 207 Mbp). This is expected, as the frequency of cut sites is dependent on the size of the genome and the properties of the sequence (e.g., GC content). Thus, the larger the genome of interest, the more RAD loci should be expected on average. Second, for a given taxon, not all RAD protocols and enzyme combinations yield the same number of loci. Enzymes with shorter recognition sequence will cut more often and yield more loci. For example, *PstI* and *SbfI* have similar GC-rich recognition sequences. However, the shorter recognition sequence in *PstI* results in a yield of loci an order of magnitude higher than *SbfI* across all genomes. GC content has similar effects. In the AT-rich genome of the *Heliconius melpomene* butterfly (GC: 32.8%), an AT-rich cutter such as *EcoRI* yields far more loci than the GC-rich *PstI*, even when their recognition sequences are both 6-bp long. Last, for a given main cutter, double-digest libraries result in fewer loci than single-digest libraries. This happens because single-digest RAD loci are defined just by the presence for the recognition sequence of the main cutter. In other words, each cut site found will always yield two loci. In contrast, in ddRADseq, the loci are defined by the presence of both enzymes. A locus will only be recovered if the recognition sequences of the primary and secondary enzymes are found within the desired insert size.

#### 2.3.2. Estimating sequencing coverage

The simulated number of RAD loci can then be used to estimate the sequencing capacity required for the experiment. For a fixed amount of sequencing (i.e., a single Illumina sequencing lane), the sequencing capacity establishes the relationship between the number of samples that can be multiplexed and the depth of coverage that will result:

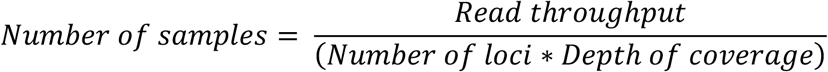

Using the number of loci simulated above for *E. maclovinus*, we can then calculate the sequencing capacity of the experiment (Table 2). Ideally, we want to sequence each sample at a 30x coverage, meaning that each individual locus is observed in 30 independent reads. On a single Illumina HiSeq 4000 lane, which yields roughly 350 million read pairs, we would be able to sequence about 222 individuals at a depth of coverage of 30x. If for example, we only had 50 individuals to sequence and we prepare a library with *SbfI*, we would obtain a coverage of 133x per sample. In this example, this high coverage would indicate that we are likely over-sequencing each sample and that, in the absence of more individuals, we could afford to use a library preparation that yields more loci.

**Table 2:** Estimates of sequencing coverage.

| Enzyme | Loci | Number of Samples at 30X | Coverage for 50 Samples |
| --- | --- | --- | --- |
| *SbfI* | 52,644 | 221.6 | 133.0 |
| *PstI* | 868,478 | 13.4 | 8.1 |
| *EcoRI* | 209,284 | 55.7 | 33.4 |
| *SbfI-MspI* | 7,939 | 1469.5 | 881.7 |
| *SbfI-MseI* | 31,135 | 374.7 | 224.8 |

### 2.4. Cleaning and demultiplexing data

After library construction and sequencing, the user will receive one or more FASTQ files (or pairs of files, if working with paired-end data) containing the multiplexed data for all individuals sequenced in the library. Before assembling loci, in most cases the raw data needs to be demultiplexed (i.e., split between the different individuals), and it should always be cleaned. We encourage users to accept only raw data from the sequencing core, as opposed to length-trimmed or quality-filtered reads. We will create an output directory to hold our unmodified, raw data we received from the sequencing core, our barcodes file, which we will create, and our cleaned/demultiplexed data that will be output by `process_radtags`:

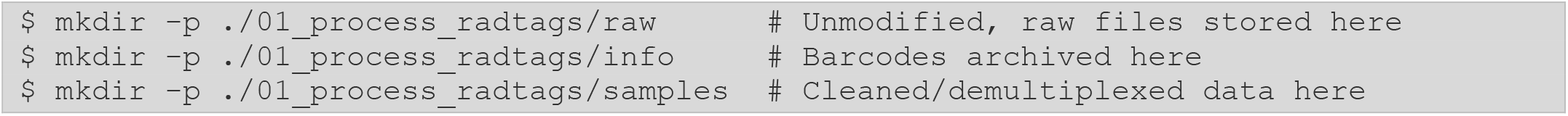

When preparing the molecular library, each individual sample was marked with a barcode, a unique molecular identifier that is present in the P1 adapter that gets ligated to the cut site. This barcode gets sequenced alongside the rest of the DNA molecule, meaning that we can then use it to assign reads to their individual of origin. We specify these identifiers (which should have been recorded when the molecular protocol was performed) in a file that we then pass to the `process_radtags` program in Stacks, containing both the sequence of the barcode and the name of the sample. In the case of our *E. maclovinus* library, after creating the file with any text-editing program (e.g., `nano`, `vi`, or `emacs`) the barcode file looks like this:

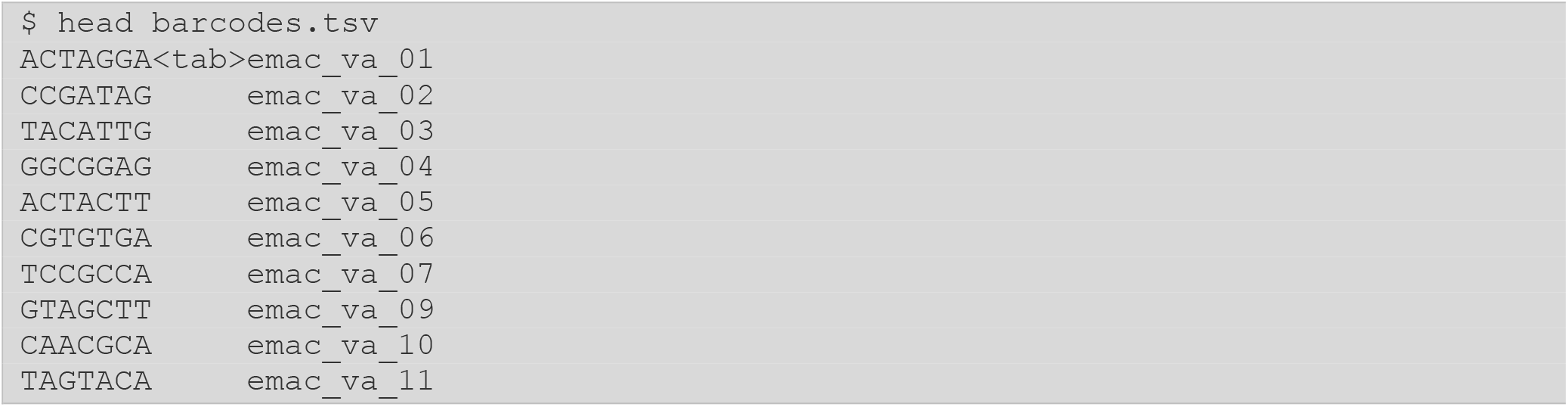

*Note! The ‘*`<tab>`*‘ text in the barcodes file is meant to signify a tab character as the file is a tab-separated file, do not include ‘*`<tab>` *‘ but rather, press the Tab key*.

*The file contains two columns separated by tabs. The first column specifies the barcode sequence and the second the sample ID. Each row corresponds to a single sample*.

*Different RAD library preparations can have different barcode configurations, including combinatorial barcodes on the single- and paried-end reads and/or incorporating the Illumina index barcodes. More information can be found in the S**tacks* *manual*.

In addition to demultiplexing, `process_radtags` can also be used to clean the data, removing reads with uncalled or low-quality bases, as well as discarding those without a recognizable barcode (since they cannot be assigned to an individual) or an intact RE remnant cut site sequence (as they might not originate from a RAD locus in the genome).

For our *E. maclovinus* dataset, we tell `process_radtags` the path to our raw data files (`--path`), as well as the path to the output directory where all the individual sample files will be saved (`--outpath`), and the location of the barcode file shown above (`--barcodes`). We also specify that our input data contains paired reads (`--paired`). To clean the data, we want to make sure that all reads with uncalled bases (`--clean`) or low-quality scores (`--quality`) are removed. Both barcodes and restriction sites will be *rescued* (`--rescue`), meaning that they will be corrected if a small number of mismatches from the expected sequence are detected. To check for the RE remnant sequence, we also specify that the library was prepared with a single RE, in this case *SbfI* (`--renz-1 sbfI`).

Our complete `process_radtags` command looks like this:

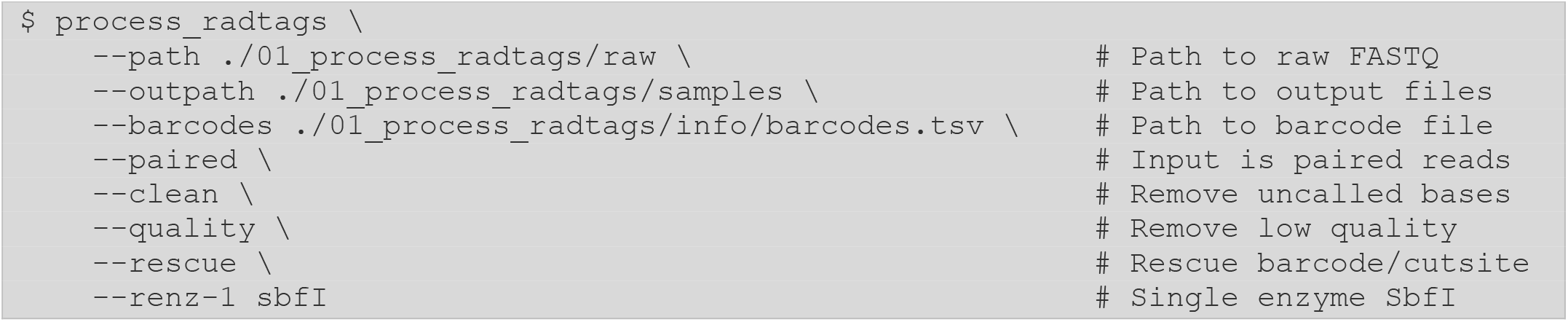

#### 2.4.1. Assaying the processing and cleaning of the raw data

Once this command is complete, a detailed breakdown of all the filtering and demultiplexing can be found in the `process_radtags.raw.log` file. From this log file, we can use the `stacks-dist-extract` utility to extract specific distributions from the different Stacks log files (Stacks log files are text files that can be viewed with any text-viewing program, using `stacks-dist-extract` is a convenience). First, we want to look at the library-wide statistics – how well did the library perform overall – examining the `total_raw_read_counts` distribution:

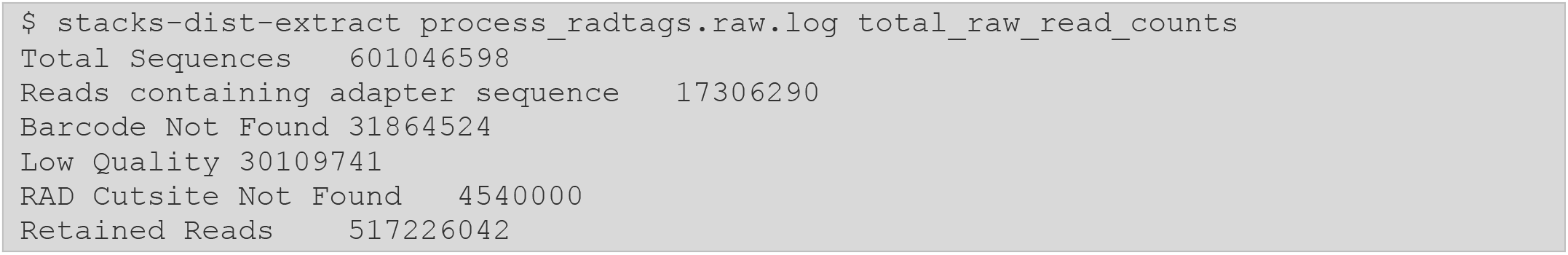

We had 601 million reads in our library, of which we retained 517 million (86%). Notice that the majority of reads lost were due to the lack of an intact barcode (31.8 million) or due to low sequencing quality (30 million). Even after these losses, we still retain a significant proportion of the sequencing and can confidently proceed with further analysis. Normally, we expect to retain between 80-90% of the total sequenced reads in the library. In scenarios were the proportion of retained reads is low, the user should assess the distribution above. Are the majority of reads lost because due to missing barcodes? If so, verify that the information in the barcode file is correct. Do many reads lack a RAD cutsite? Perhaps the enzyme passed to `process_radtags` does not match the library preparation, or the specific RAD P1 adapter used in the protocol may have included extra nucleotides in the barcode/RE cut site region. Lastly, are many reads lost due to low quality? Here, the user should contact the sequencing facility to assess any problems with the sequencing lane.

Once the library-wide assessment is performed, the next step is to verify the proportion of reads kept between the different samples. We can again use the `stacks-dist-extract` program to extract the `per_barcode_raw_read_counts` distribution from the log.

Looking at the first ten lines of the distribution we can see that it counts the number of total reads in the library for each barcode (sample), as well as the number of reads retained and lost due to the lack of a restriction enzyme cut site or low sequencing quality.

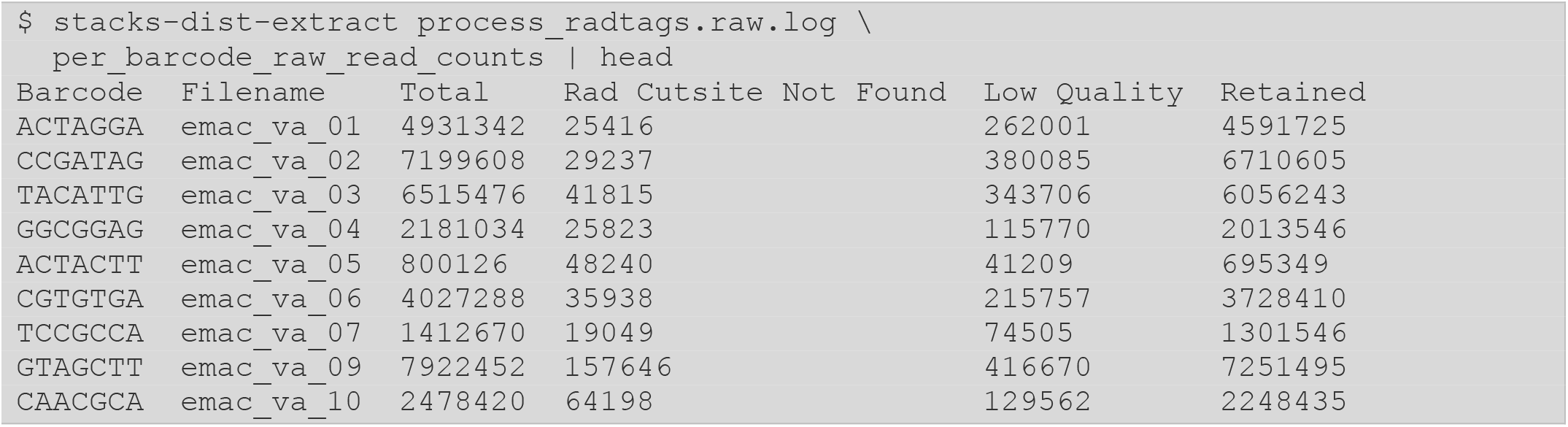

The user can capture this data into a file to load into R (R Core Team 2021) or to another statistical software and calculate detailed distributions of the proportion of reads kept and lost for each sample. Regardless of the method used, the user should check for at least three things: 1) the proportion of each sample in the total library, as well as 2) the number and 3) proportion of reads retained in each sample. The Appendix links to an R script providing an example on how to perform these calculations.

The proportion of reads per sample (Fig. 1A) roughly represents the proportion at which individuals were multiplexed in the library. For example, here we can observe that some samples composed over 3.5% of the library, while others were present in less than 0.04%. Such differences are expected given a combination of effects during library preparation (e.g., human error, DNA quality, equipment sensitivity). However, the user should monitor for larger effects, such as biases between populations (Fig. 1B) or sequencing lanes, or for cases in which one or a few samples dominate the sequencing as this may imply that library contamination occurred.

**Figure 1.**
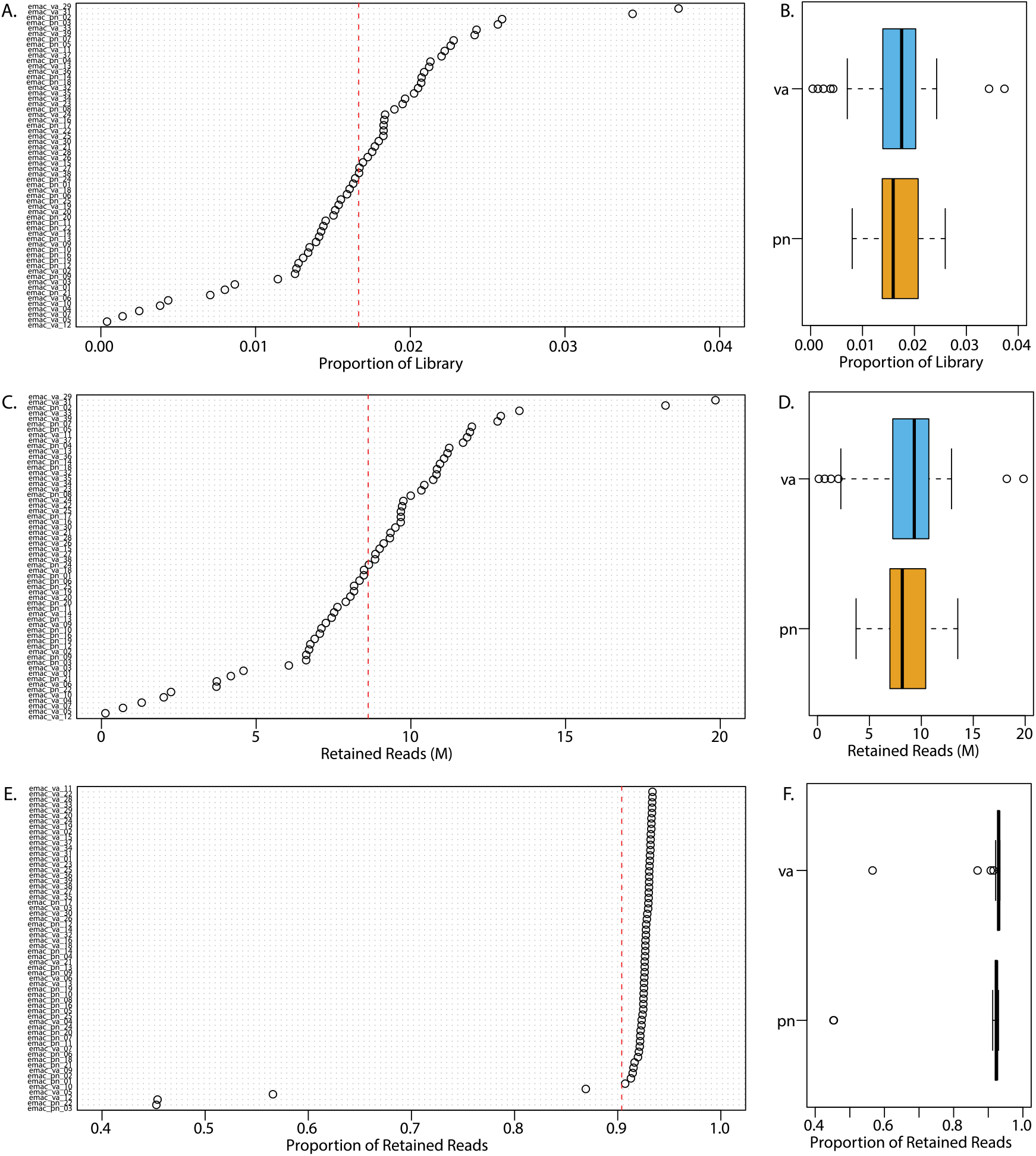
Distribution of reads per sample after process_radtags. (A) Proportion of raw sample reads in the total library (sum of raw reads across all samples). Red dashed line shows the mean proportion of samples reads in the total library. (B) Proportion of total library between Valdivia (`va`, blue) and Puerto Natales (`pn`, yellow) samples. (C) Number of retained reads per sample, in millions of reads. (D) Number of retained reads between populations. (E) Proportion of retained reads per sample. (F) Proportion of retained reads between populations.

Next, the user should check the number of reads retained per sample, as it is an early metric to assess sequencing coverage (Fig. 1C). In our library, we can observe variation between samples, ranging from 130k to 19 million reads. This variation is expected given the differences in per-sample library composition described above. Similarly, while there are some small differences in the expected coverage between populations (Fig. 1D), they do not indicate any large-scale biases in library preparation. Note that, given the very low number of retained reads for some samples, it is a good practice to remove these individuals from further analyses. In the case of this library, we will remove the two samples retaining less than 1 million reads (`emac_va_05` and `emac_va_12`).

*Note! This threshold of 1 million reads used to remove samples showing early low coverage is not an absolute one. It is used here based on the properties of the dataset used and could vary depending on the number of markers recovered from the genome and the overall number of reads per sample*.

The last check is the percentage of reads retained per sample. While we originally had observed that we had retained 86% of all reads in the library, this original assessment did not account for variation between samples. As we can observe from Figure 1E, a subset of the samples retained only a small fraction of the total reads and should be assessed in more detail.

### 2.5. *De novo* Analysis

When a reference genome is not available, is of low or questionable quality (e.g., low N50, low contiguity, certain regions missing), or when a researcher desires to conduct an analysis independent from a particular assembly (e.g., the construction of a genetic map), then a *de novo* analysis is the recommended approach. When a high quality, complete reference is available, it will typically yield superior results, but the option to assemble *de novo* provides the researcher with a lot of flexibility and the ability to test hypotheses independent of the genome. Here we will show the major steps involved, starting with parameter optimization, followed by the execution of the core pipeline, and following up with a set of methods to evaluate the success of the assembly.

#### 2.5.1. Parameter Optimization

When analyzing RADseq data without a reference genome, it is important to optimize the parameters that will define the *de novo* assembly of loci. There are two main parameters in Stacks that control the assembly of alleles into loci within individuals and the matching of these loci, in a catalog, between individuals. The initial assembly of loci within each sample happens in `ustacks`, where the parameter `-M` specifies the number of mismatches allowed when matching stacks, i.e., putative alleles, into loci. Setting this parameter too low results in under-merging – leaving two alleles as separate, homozygous loci. In contrast, if `-M` is too high alleles can be over-merged, collapsing alleles of different loci into a single locus. The value is `-M` is roughly equivalent to the level of biological polymorphism present in an individual given a particular read length (e.g., `-M` should be set lower when using very short reads).

Once loci are assembled within samples, they are then merged by `cstacks` into a catalog of loci, representing the metapopulation. The `-n` parameter specifies the number of mismatches between the loci of individuals and the catalog, playing the same role as `-M` in `ustacks`. For example, a variable locus may have been assembled as a homozygous locus in one individual, and as a homozygous locus with the alternative allele in a second individual. The `-n` parameter allows `cstacks` to merge these loci into a proper heterozygous locus for the population with the same side effects if `-n` is set too low (under-merging) or too high (over-merging). The value of `-n` is roughly equivalent to the level of biological polymorphism present across the population. If working with several populations, `-n` should also account the genetic distance between them (i.e., `-n` might need to be larger if the populations are divergent or are separate species).

Several protocols have been published regarding the optimization of assembly parameters for *de novo* analysis (Paris et al. 2017; Rochette and Catchen 2017; McCartney-Melstad et al. 2019; Heller et al. 2021); McCartney-Melstad et al. (2019) used a method based on the relation of sequence similarity and the fraction of sequences flagged paralogous, while Heller et al. (2021) performed a *de novo* catalog assembly with default parameters on a subset of individuals, after which reads for all samples are aligned to the catalog consensus sequences and analyzed as whole-genome resequencing data. In this tutorial, we will be focusing on the R80 method first proposed by Paris et al. (2017) and described further by Rochette and Catchen (2017). Briefly, this method looks for the assembly parameters that maximize the number of R80 loci in the dataset (i.e., the number of polymorphic loci present in at least 80% of the samples). Values of `-M` and `-n` outside of this optimum result in the over- and under-merging of loci, leading to an increased proportion of missing data across the dataset.

To perform the parameter optimization using the R80 method, we will run the core pipeline several times on a subset of individuals, iterating over increasing values of `-M` and `-n` in each run. In the first run, both `ustacks -M` and `cstacks -n` will be set to ‘`1` ’ — allowing for one mismatch between alleles in both the individuals and the population. These values will then be increased until a maximum of 10-12 mismatches is reached. Once the iterations are complete, the user will calculate the change in R80 loci between subsequent runs (i.e., did the number of R80 loci increase or decrease after the assembly values were increased by one). When optimum assembly parameters are found, the change in the number of R80 loci will approach zero and remain positive. Exceeding the optimal value will result in a negative number, indicating a loss of R80 loci.

Before we can optimize, we must select a subset of samples on which to run the analysis. These samples should be representative of the whole library, so it is ideal to include samples across all studied populations or groups. Similarly, it is best to include samples with average amounts of input data (which can be assessed from the method described in Section 2.4.1). For the *E. maclovinus* library, we selected 20 individuals for parameter optimization, 10 each from the Valdivia and Puerto Natales populations. Each sample contained between 7.6 to 10.9 million reads (average of 8.9 million). We will record the prefix sample names (i.e., the filename for the sample without any suffixes such as “`.1.fq.gz` ”) of these samples in a *population map* text file (`popmap.tsv)`, a list associating individuals with a population grouping. Here we have a single ‘population’, and we arbitrarily refer to it as ‘`opt`’, to signify *optimization*.

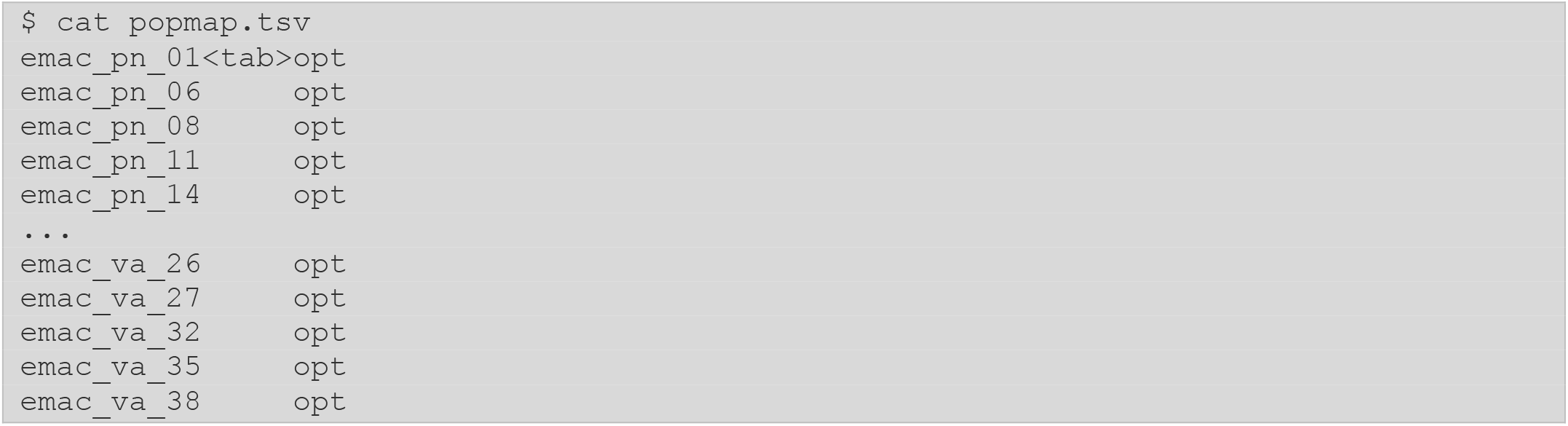

After creating the population map, the next step is to create an output directory for the current run of the pipeline. For example, here we make a new directory using the current value of `-M` as part of the name.

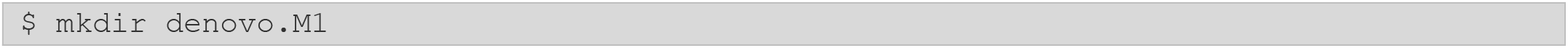

With the population map and output directory, we can now prepare the Stacks `denovo_map.pl` program (`denovo_map.pl` is a Perl wrapper script that will execute the core pipeline for us). First, we specify the path to our input reads (`--samples`), the path to the population map file (`--popmap`), and the path to our output directory (`--out-path`). In the case of the *E. maclovinus* library, we have paired-end reads and thus want them included in the analysis (`--paired`). We also want to remove PCR duplicate reads (`--rm-pcr-duplicates`), as well as only output R80 loci (`--min-samples-per-pop 0.8`). Last, we set our values if `-M` and `-n` to our current iteration of the pipeline, in this case ‘`1`’.

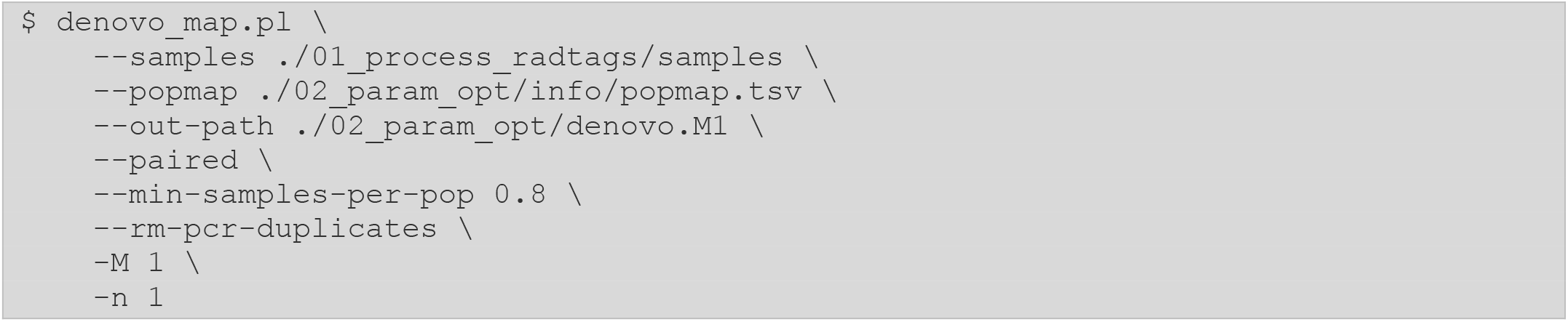

The code above runs the pipeline once on our 20 samples, using the specified assembly parameter values. The iteration across all desired values can be automated using a shell loop, and we provide an example script, listed in the Appendix.

Once Stacks is complete, the number of polymorphic R80 loci can be extracted from the output of `populations`. In this case, we are using the `population.sumstats.tsv` file, which contains the summary statistics for each variant site per population in the dataset (e.g., sample counts, allele frequencies, observed and expected heterozygosity, etc.). The first column of this file contains the locus ID from which each variant site originates, and performing a tally of these unique values would provide us with the number of R80 polymorphic loci:

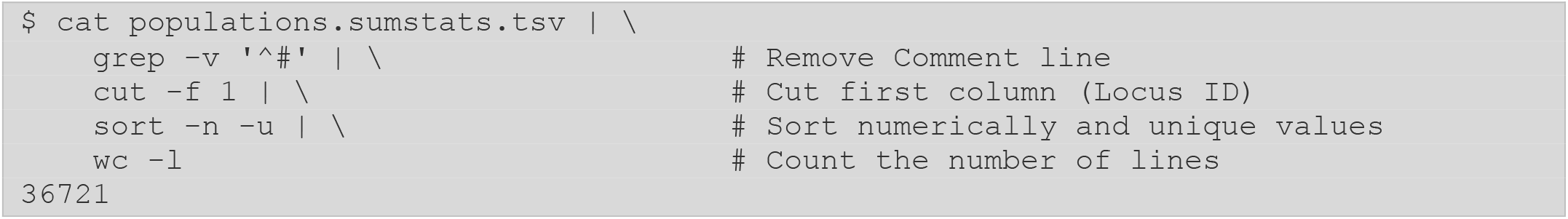

For this dataset, allowing for 1 mismatch between the alleles of individuals and across the population provides 36,721 R80 loci. This value is similar to that reported in the `population.log` file, except that the value in the log reports the total number of polymorphic and invariant R80 loci (37,576).

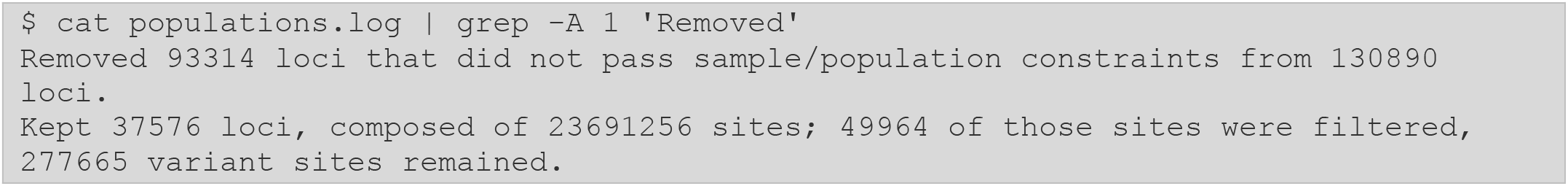

Once the core *de novo* pipeline has been run for all `-M`/`-n` values, we can proceed to calculate the change in R80 loci between runs (Table 3). To do this, we can subtract the number of R80 loci found in a run, with the number of R80 loci found in the previous run (i.e., `M=1` to `M=2`, `M=2` to `M=3`, etc.). Plotting these results, we look for the value in which the change in R80 loci approaches zero, while remaining positive (Fig. 2). Here, increasing `-M`/`-n` from 2 to 3 results in a gain of 121 R80 loci, while increasing from 3 to 4 results in a loss of 40 loci. This indicates that an `-M`/`-n` of 3 is the optimal value for *de novo* loci assembly for the *E. maclovinus* dataset.

**Table 3:** Change in R80 loci.

| $M/n$ | Number of R80 Loci | Change in R80 Loci |
| --- | --- | --- |
| 1 | 36,721 | - |
| 2 | 37,647 | 926 |
| 3 | 37,768 | 121 |
| 4 | 37,728 | -40 |
| 5 | 37,589 | -139 |
| 6 | 37,518 | -71 |
| 7 | 37,449 | -69 |
| 8 | 37,430 | -19 |
| 9 | 37,301 | -129 |
| 10 | 37,244 | -57 |
| 11 | 37,190 | -54 |
| 12 | 37,066 | -124 |

**Figure 2.**
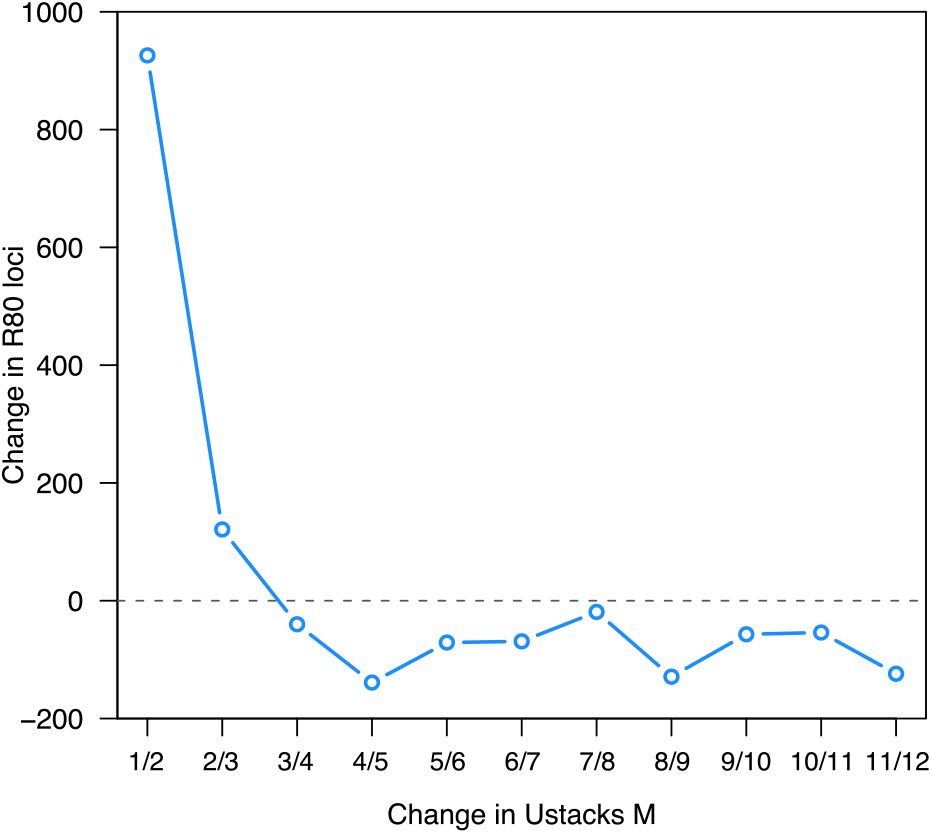
Change in R80 loci during *de novo* parameter optimization. Blue line shows the change in loci kept in 80% of individuals with the successive increase of `ustacks -M`. The optimal assembly parameter value for the *E. maclovinus* dataset is an `-M` of 3.

*Note! The optimization procedure will not always converge for all datasets. It is possible that there are several, equivalent values for* `-M` *and* `-n` *in a dataset, as well as scenarios in which* `-M` *and* `-n` *are not equal. Additionally, when working with several divergent populations, the optimal parameters might even be different for each population. Other factors can also affect parameter optimization, such as the presence of repetitive loci in the genome and highly polymorphic indels in the metapopulation. We encourage users to explore these parameters, but we advise against infinitely iterating over all possible combinations in order to find perfect parameters. The maximization of R80 loci is just the metric of the optimization process, not the underlying biological signal present in the data*.

#### 2.5.2. Core *de novo* pipeline execution

With the completion of parameter optimization, we are ready to employ those parameters to assemble our RAD loci and genotype all samples. Recall that R80 loci are simply a target we employed for optimization. Now, we will deal with the full dataset, which of course includes loci present in a range of sample numbers, from as few as one to all samples. We will have the option to filter these data using `populations` (Section 2.8), once the core pipeline is complete.

We will prepare a new population map that includes all samples and now lists our biologically meaningful population associations (`va` and `pn`, here based on geography). Note that based on the analysis of our `process_radtags` results, we have removed the two samples with less than 1M reads, as including them will skew our filtering results:

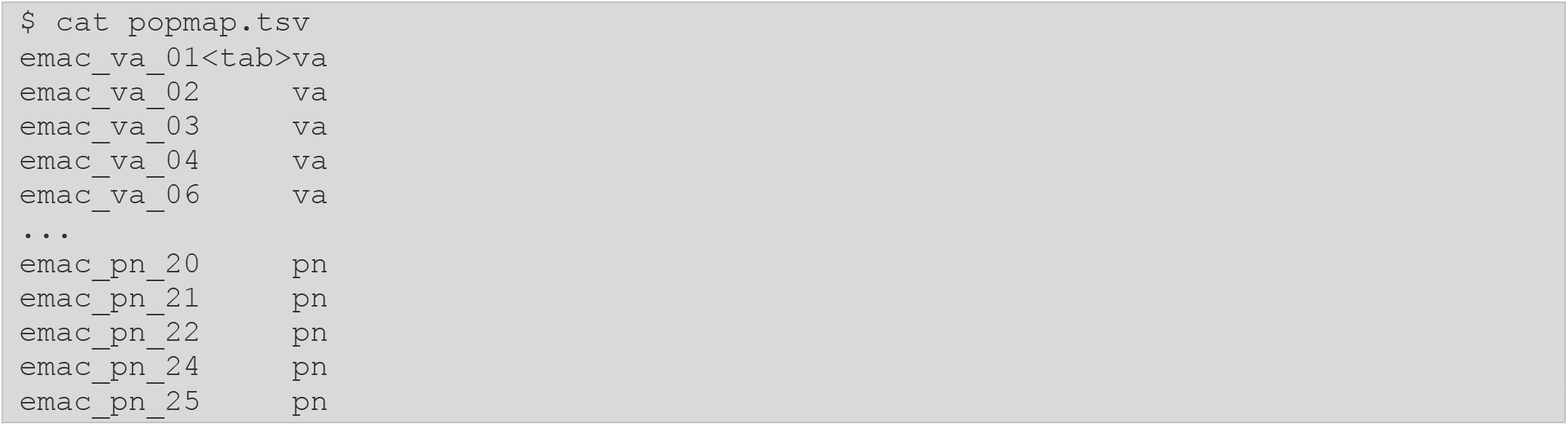

We make a new output directory:

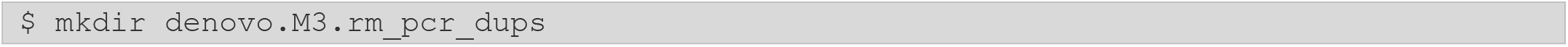

And execute the `denovo_map.pl` wrapper, which will run Stacks *de novo* component programs, removing the PCR duplicates found from our paired-end reads:

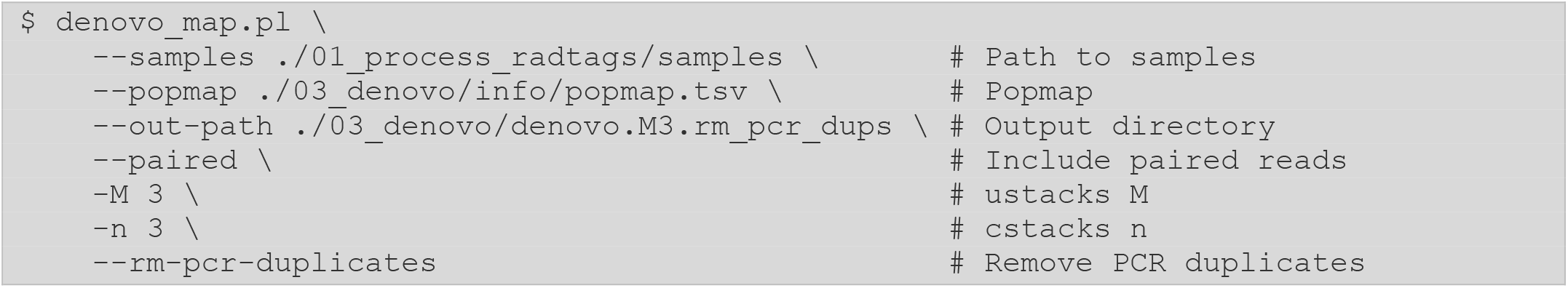

#### 2.5.3. Assaying the effectiveness of *de novo* assembly

Once the pipeline is finished, we will focus on three aspects of the analysis to assay our results: sequencing coverage, number of PCR duplicates, and phasing of SNPs into haplotypes within each fully-assembled (single- and paired-end) locus. Sequencing coverage can be assessed at different stages of a RAD analysis (n.b., we initially assessed coverage after demultiplexing the raw, unassembled reads in Section 2.4.1). The first stage during the core pipeline run is following the execution of `ustacks`, in which the single-end reads from each sample are processed independently and prior to the removal of PCR duplicates. We can check these results by extracting the per-sample coverage distribution from the `denovo_map.log` file using the `stacks-dist-extract` utility:

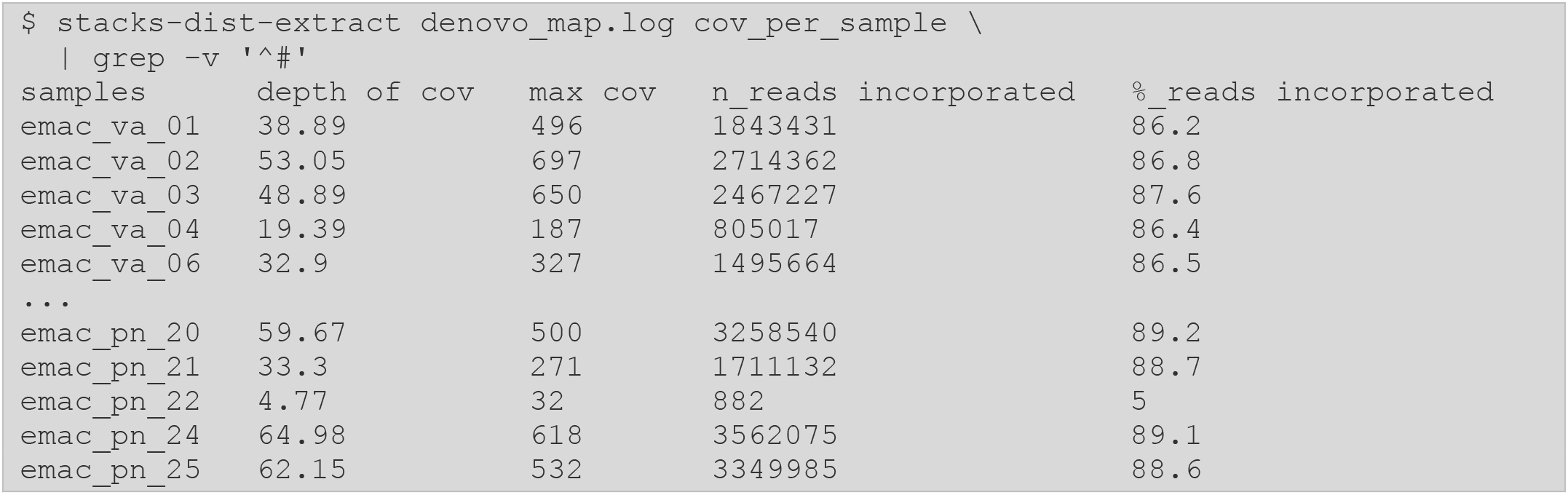

The distribution demonstrates a high variation in coverage between samples, which matches the distribution of reads observed after library demultiplexing. While average coverage is very high, a few samples are outliers. For example, `emac_pn_22` contained only a small number of forward reads (∼17k) of which only 882 (5%) were incorporated into loci by `ustacks`.

While the final assessment of coverage, accounting for PCR duplicates, will take place after running `gstacks`, this early assessment is important as it might highlight problems with some of the s amples. Trends like these, in which only a small fraction of reads were used in `ustacks`, can imply problems in library preparation, such as poor DNA quality and even the presence of contamination.

After `gstacks` executes, building the paired-end contig and incorporating it into the single-end locus, we can verify coverage once again, as well as checking the levels of PCR duplicates and rates of phasing. A summary of these metrics for the whole library can be found in the `gstacks.log` file, as well as information on the locus assembly and genotyping:

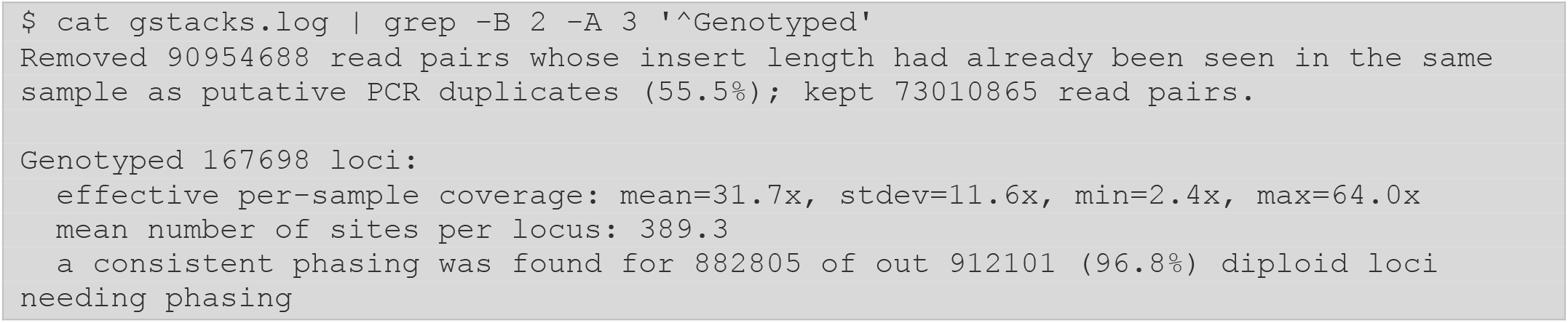

We first see that the final coverage is 31.7x, almost half of the coverage observed in `ustacks`. This is because this final coverage is calculated after the removal of PCR duplicate reads, which account for 55.5% of reads in this library.

*Note! For single-digest RAD libraries, PCR duplicates are determined by looking at each pair of reads in a particular locus and if the paired-end aligns to the same position as another pair of reads, it is assumed to be a duplicate produced during PCR amplification*.

Our post-duplicate levels of coverage are still robust and will yield confident genotypes. The high number of PCR duplicates in this library likely comes as a function of the very high sequence coverage. At this point, if coverage has dropped significantly below 10x, the researcher should consider that genotype calls may be of low confidence and biased toward homozygotes, and data may be widely missing from loci throughout the population (see Section 2.9.3).

Last, we can check how well individual genotypes could be phased into two distinct haplotypes (diploid alleles) in each individual at each locus. The `gstacks` program can phase SNPs into haplotypes by linking alleles together that occur in the same set of paired (and linked) reads. In *E. maclovinus*, 96.8% of all loci could be phased. The proportion of unphased loci often represent the presence of repetitive and paralogous sequences in the genome, though typically low phasing demonstrates problems in the assembly of loci (for example, excessive over-merging).

A further breakdown of the results from `gstacks` can be extracted from the `gstacks.log.distribs` file using the `stacks-dist-extract` utility. It provides information on the number of raw loci present in each sample, as well as coverage, and PCR duplicate rates. Coverage is reported both as general average coverage (`mean_cov`) and the average coverage adjusted by the number of samples present at a locus (`mean_cov_ns`). We will use the adjusted mean to assess the coverage across samples in this section.

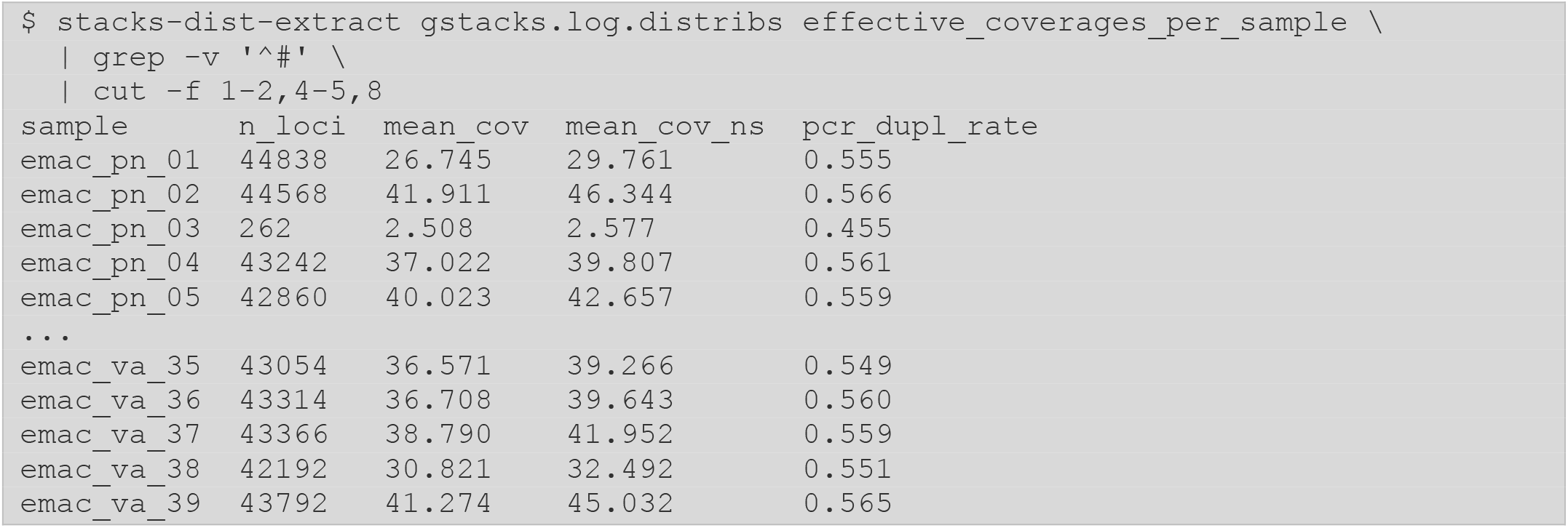

*Note! We are using the* `cut` *command to show only some of the columns in this file for readability purposes*.

If we save this distribution to a file, we can use R to calculate further statistics on the coverage and PCR duplicate data. In particular, we want to observe the distribution of coverage across the samples (Fig. 3A), so we can determine if any additional samples need to be removed from the analysis (See the Appendix for a link to the R code). In accordance with the results from `ustacks`, we observe a subset of samples (e.g., `emac_pn_03` and `emac_pn_22`) with very low coverage (<5x), which will be removed from downstream analyses. The same samples also display a reduced proportion of PCR duplicates (Fig. 3C), but this is likely a function of their reduced coverage. In addition, we can also verify if there are any biases between the populations of groups in the library. While we, again, observe differences between the `pn` and `va` populations, we do not see any large-scale biases in coverage (Fig. 3B) or PCR duplicate rate (Fig. 3D).

**Figure 3.**
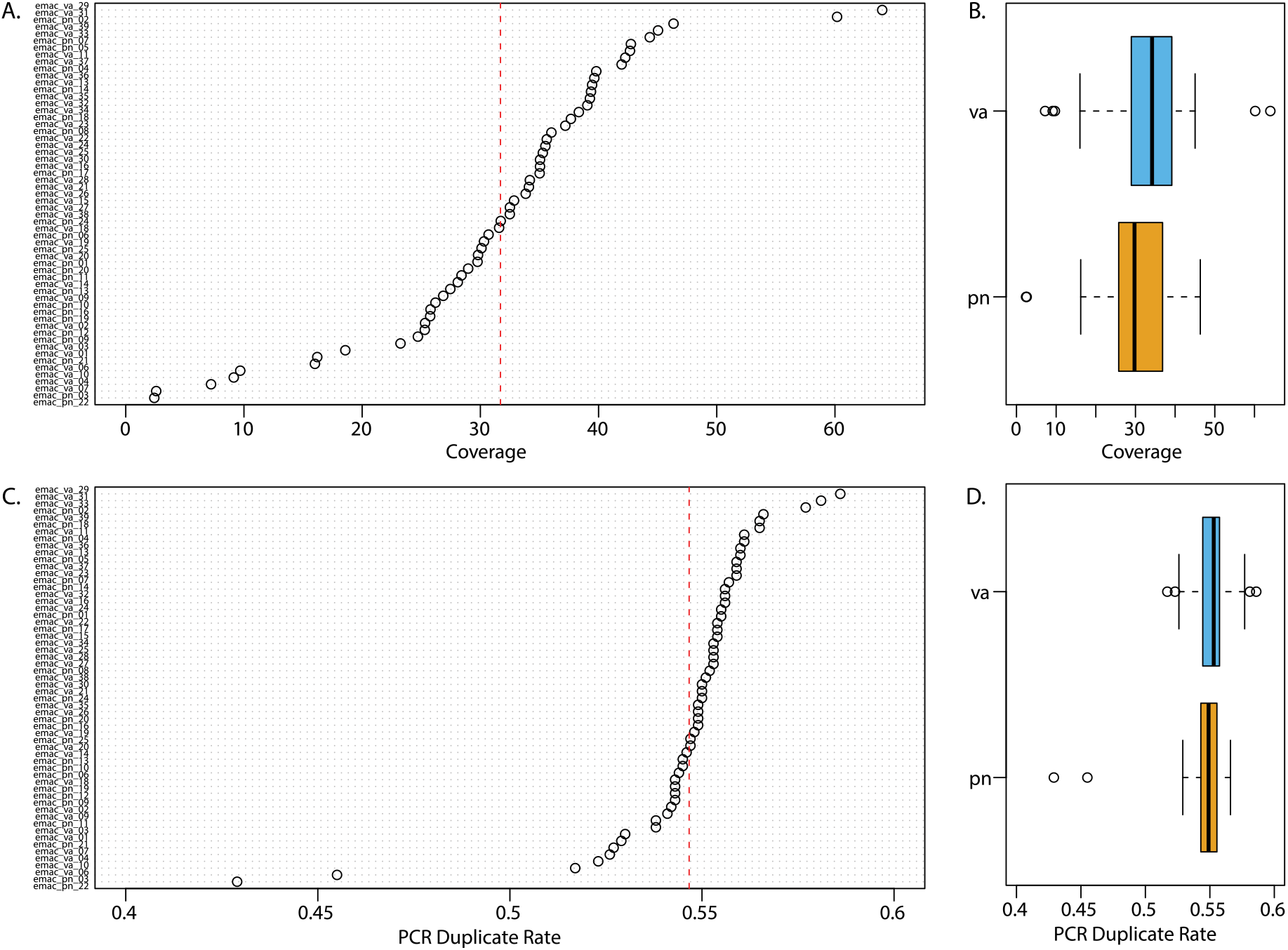
Sequencing coverage and PCR duplicate statistics. (A) Non-redundant depth of coverage per sample. Red dashed line shows the mean coverage across all samples. (B) Non-redundant depth of coverage between Valdivia (`va`, blue) and Puerto Natales (`pn`, yellow) populations. (C). PCR duplicate rate, proportion of reads identified as duplicates, per sample. (D). PCR duplicate rate between populations.

We can obtain additional information by extracting per-sample phasing statistics. From this distribution we can look at both the number of genotypes called per individual, as well as the proportion of loci that could not be phased. These metrics can also be used to identify any problematic samples that should be removed from the analysis. In this case the same two samples with low coverage (`emac_pn_03` and `emac_pn_22`) also display outlier patterns of phasing and number of genotypes.

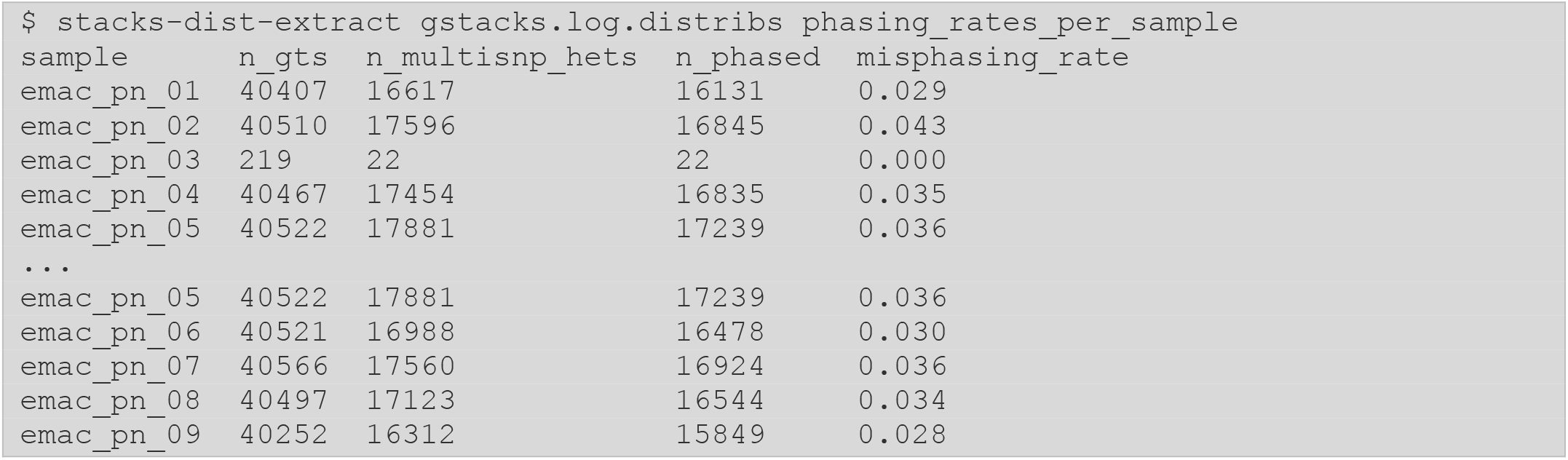

From here, the core pipeline run is complete. While the outlier samples we identified have contributed some data to the metapopulation catalog, we can prevent them from being included in our downstream analysis simply by removing them from the population map. If we want to ensure that no data at all was included from the outliers, we would re-execute `denovo_map.pl` after adjusting the population map.

We will next show how to complete an analogous core analysis using a reference genome, followed by a hybrid *de novo*/reference assembly method called *integration*. After that, the three methods converge when we will apply a population genetics frame to the processed data using the `populations` program.

### 2.6. Reference-aligned Analysis

When a reference genome is available for the species of interest, Stacks can be run in reference mode. A local assembly of loci is still performed, using the position of the aligned reads in the genome to identify the reads belonging to a locus, in contrast of the clustering based on sequencing similarity used by the *de novo* approach. Thus, no parameter optimization is required. However, note that locus assembly is performed independent of the reference genome, as the reference is not used as a template or as a source of alleles for SNP calling. A reference-aligned analysis allows us to place the genotypes and the population-level metrics in the context of the chromosomes, enabling analyses such as genome scans of genetic divergence (Bassham et al. 2018), measuring linkage disequilibrium and identifying structural variants (Chen et al. 2019; Mérot et al. 2021), and fine-scale coalescent reconstruction along chromosomes (Nelson and Cresko 2018).

Before we can run Stacks, we need to align the reads generated by `process_radtags`(Section 2.4.1) to the reference of interest. This can be done using any standard short-read aligner, we will be using BWA (Li 2013) here. Alignments will then be sorted using SAMTOOLS (Li et al. 2009). We will create a set of directories to hold the reference genome and the aligned samples:

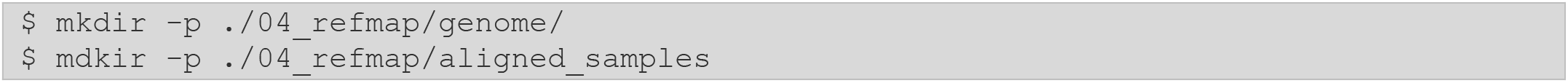

Before we can align, we need to generate a searchable database of the reference genome that can be accessed by the aligner. For our *E. maclovinus* reference, we will generate this database using the `index` mode of BWA, and we will name the resulting database as `emac_db`:

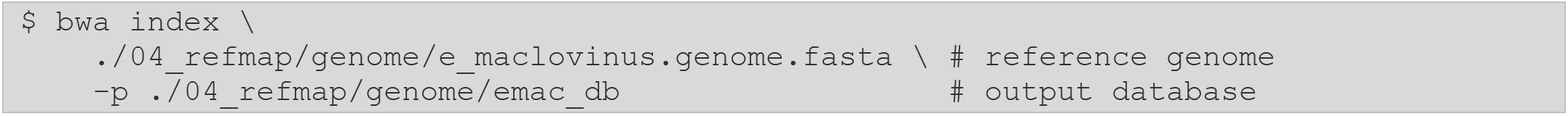

Once the database is generated, we will align each sample independently using BWA in `mem` mode. The output of BWA will then be directly piped into `samtools view`, to compress the alignments (`-b`) while preserving the headers (`-h`), and these data are likewise piped into `samtools sort`. Aside from sorting, no further processing or filtering is done.

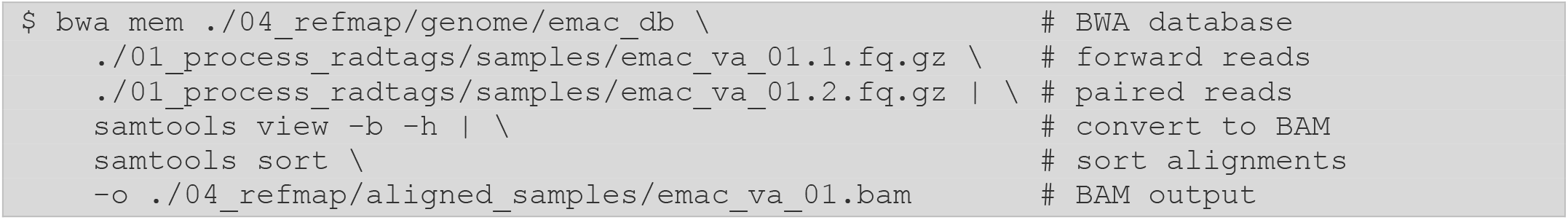

The alignment of all samples can be automated using a shell script, an example of which is linked in the Appendix, or they could be run as independent jobs on a cluster, to speed up execution.

Next, we will create the output directory for the Stacks reference run:

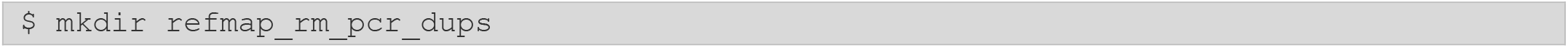

We can now prepare the command for the `ref_map.pl` program. Similar to the *de novo* wrapper, `ref_map.pl` is a Perl wrapper script used to execute the core reference-based pipeline. We will specify the path to our processed reads (`--samples`), the path to our new output directory (`--out-path`), and the path to our population map (`--popmap`). This population map file is the same used in Section 2.5.2, which classifies the samples between the `va`and `pn` populations and excludes the two individuals with low read counts. Last, we tell `gstacks` to remove PCR duplicates (`--rm-pcr-duplicates`).

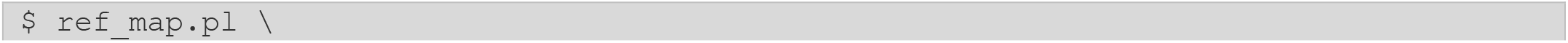
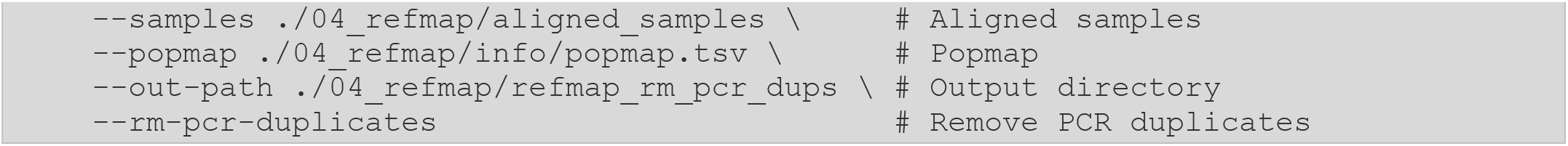

#### 2.6.1. Assaying the effectiveness of a reference-based assembly

With the completion of pipeline execution, we first want to assess our input data, particularly the quality of the alignments. The `gstacks.log`file contains information about the input alignments:

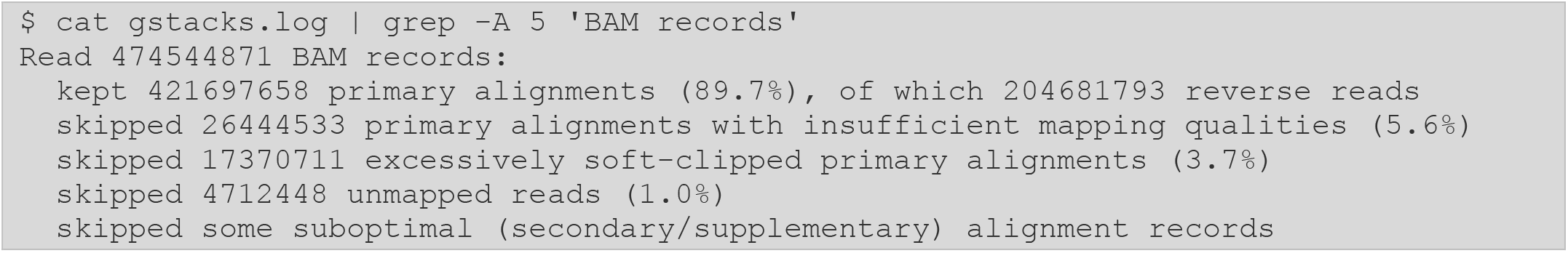

Users should verify the proportion of retained alignments and examine the main reasons alignments were discarded. While the proportion of alignments kept in this dataset is high (89.7%), users should check for the fractions of alignments discarded due to low mapping quality or excessive soft-clipping, as they might imply problems with the underlying reference, presence of non-target DNA in the library, or, in the case of using a heterospecific reference, a large evolutionary distance between the reference the species sequenced.

A quick way of visually assessing the alignments is by looking at the CIGAR string (Li et al. 2009) present in column six of the SAM/BAM file format. We can extract some of these CIGAR strings using the following command:

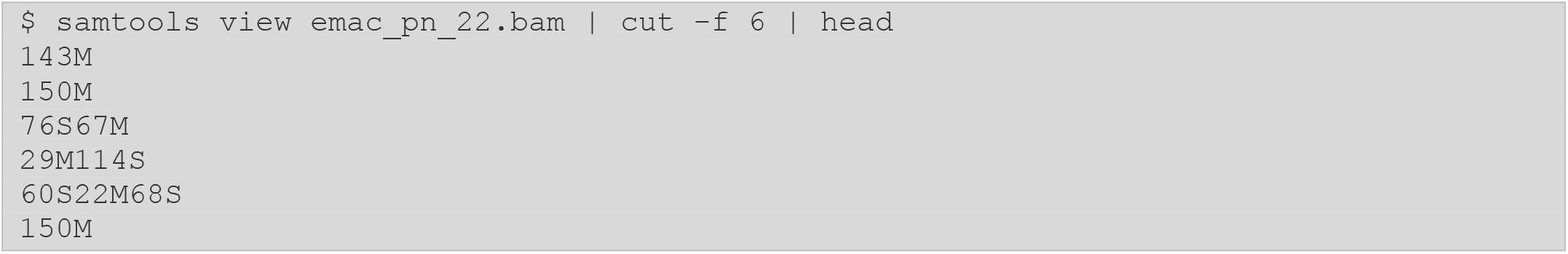

*Note! Remember that BAMs are binary and need to be processed and translated to text using the* `samtools view` *command*.

The CIGAR strings report information on the structure of the alignment. If the reads are 150bp long, a CIGAR string of 150M would present a one-to-one alignment of 150 “matches” between the reference and query sequence (n.b., *matches* implies two nucleotides are *aligned*, not necessarily identical). The presence of indels in an alignment are recorded with “`I`” and “`D`” designations (e.g., `20M1I129M` represents an alignment with 20 matches, a 1-bp insertion in the subject, and then 129 additional matches). CIGAR strings can also be used to represent partial alignments, in which a large portion of the subject read does not match the reference, represented by the “`S`” and “`H`” operations, in which part of the subject is excluded from the alignment (e.g., given `76S67M`, the first 76 bases of the reads do not align to the reference and are soft-clipped). The presence of these partial, soft-clipped alignments is expected; however, excessive soft-clipping might be a sign of a poor-quality reference, too much genetic distance between the subject species and reference, and/or improper processing of the raw reads. Moreover, the presence of soft-clipped alignments is not accounted for by the mapping quality (`MAPQ`) calculation present in the BAM file, meaning that filtering for quality alone might not remove all the partially aligned reads.

For reference-aligned data, a per-sample distribution of alignment statistics can be obtained from the `gstacks.log.distribs`file using the `stacks-dist-extract` utility:

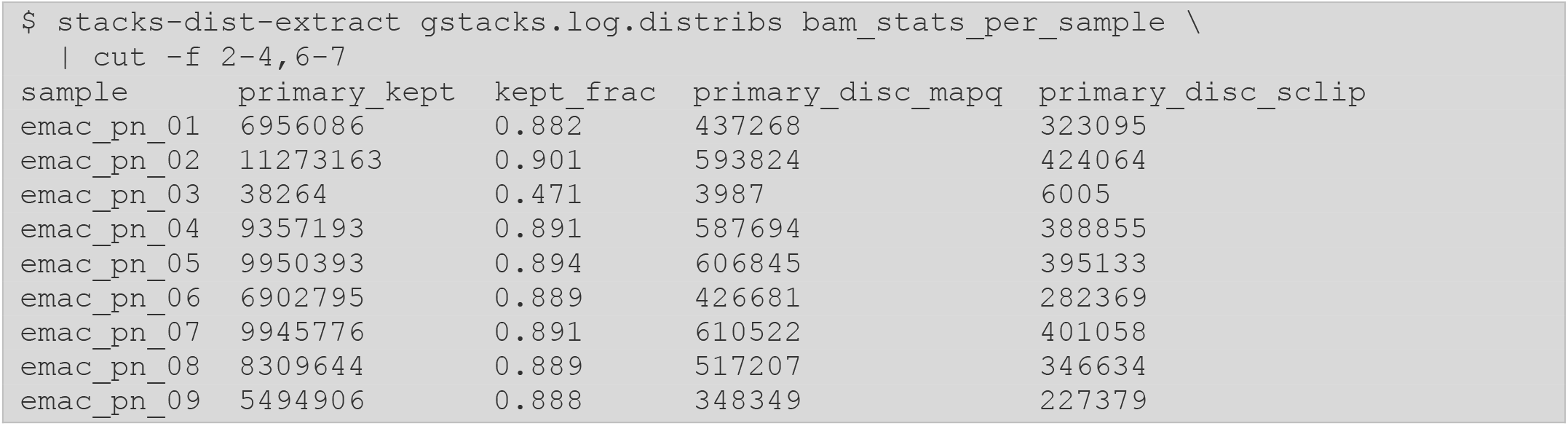

It is useful to verify two at least two alignment statistics including, 1) the number of primary records (alignments) kept per sample, and 2) the fraction of records kept. We observed that an average of 7.3M records were kept per sample, although variation is seen between the samples (Fig. 4A) — which is expected given the differences in the number of input reads and resulting coverage. When we compare the number of records kept between the two populations, we observe that, while the `va` population keeps more records overall, no large biases are observed between the two groups (Fig. 4B).

**Figure 4.**
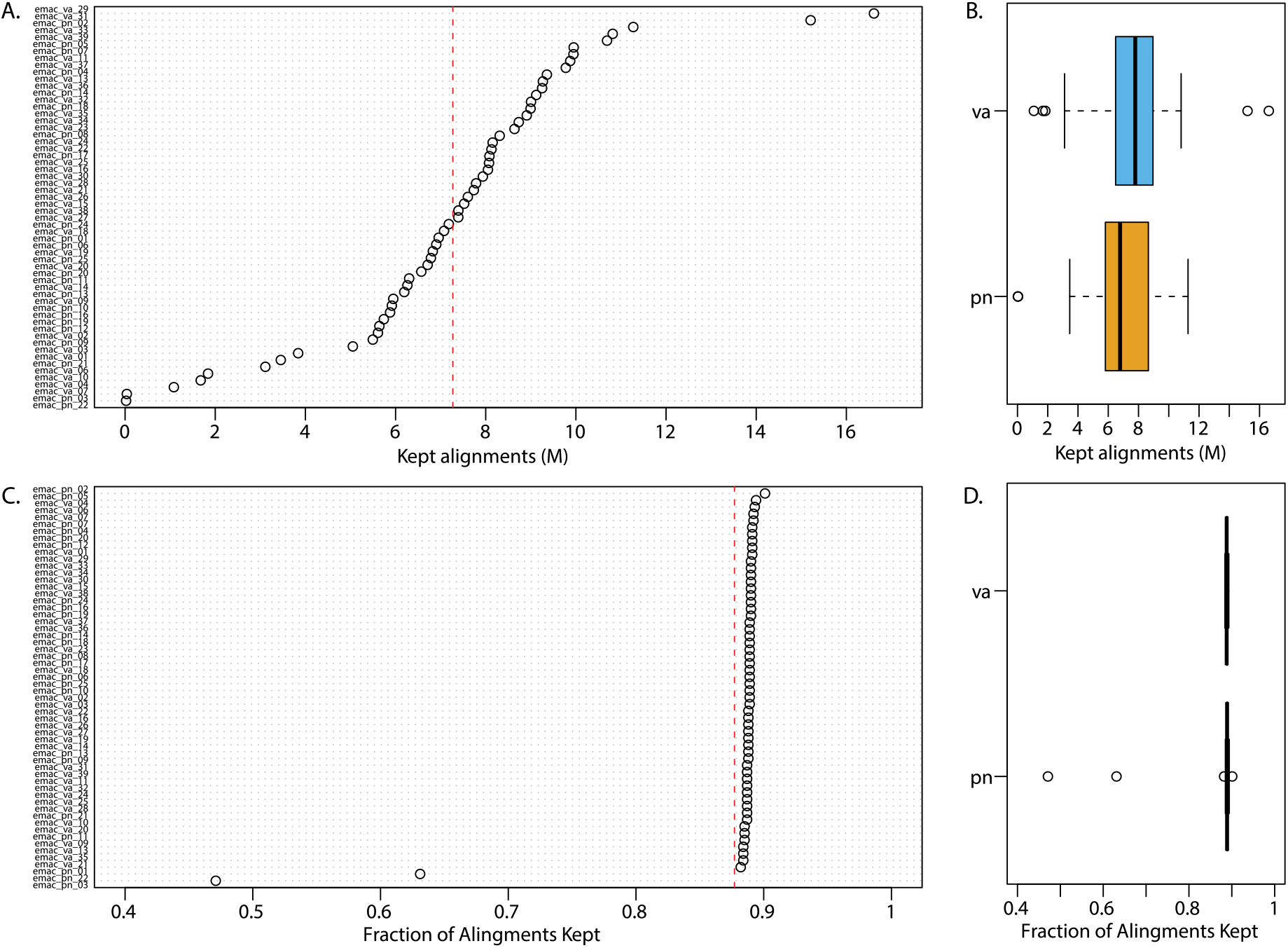
Reference-based aligned statistics. (A). Number of alignments per-sample kept by `gstacks`, in millions of alignments. Red dashed line shows the mean number of kept alignments per sample. (B) Number of kept alignments between the Valdivia (`va`, blue) and Puerto Natales (`pn`, yellow) populations. (C). Fraction of alignments kept per sample, out of the total number of reads. (D) Fraction of kept alignments between populations.

While the library has a high average proportion of kept alignments, there are some samples displaying outlier proportions of kept reads (Fig. 4C). These samples (e.g., `emac_pn_03` and `emac_pn_22`) also display very reduced coverage and can be removed from the population map for subsequent analyses. Aside from this section regarding alignments, the output of `gstacks` is otherwise identical to the one obtained in *de novo* mode. Therefore, coverage, PCR duplicates, and phasing can be assessed as described in Section 2.5.3.

### 2.7. Integrated Analysis

A high-quality reference genome (with a large N50 and chromosome-level contiguity) provides the best foundation for a RAD-based population genetics analysis. However, a *de novo* analysis can still reveal a lot about the underlying genetics of a natural population, particularly with respect to population structure, levels of heterozygosity, and coalescent and phylogenetic inferences. Stacks provides a third option: conducting a *de novo* assembly, followed by alignment of those loci to a heterospecific reference genome. This *integrated* method allows a researcher to assemble loci without a distantly related or low-quality reference affecting the results, but still providing information with respect to the colocalization of loci along the chromosomes (e.g., to visualize runs of divergence). The `stacks-integrate-alignments` program provides the ability to inject locus alignment coordinates into the Stacks catalog and related files, while providing filters to eliminate spurious alignments.

The procedure starts by assembling loci *de novo* (Section 2.5.2), and then aligning the loci in the catalog (i.e., `catalog.fa.gz`) against a reference genome of interest, obtaining a BAM output file of those alignments. Next, the `stacks-integrate-alignments` Python program is run, feeding it the *de novo* output directory, the aligned loci BAM, and specifying any filters, which can be used to require a certain percentage of the length of a locus to be aligned, as well as a minimum percent identity for an underlying alignment. These filters prevent spurious matches between loci and, e.g., repetitive elements in the heterospecific reference.

To demonstrate this functionality, we will align our *E. maclovinus de novo* loci against another non-Antarctic notothenioid, the Channel Bull Blenny, *Cottoperca gobio* (NCBI accession GCA_900634415.1). This species is in a different family from *E. maclovinus*, so we expect only a fraction of our *de novo* loci will be alignable, none the less, it is a high-quality, chromosome-level, assembly with an N50 > 25Mbp.

We create a series of directories to hold the integrated data:

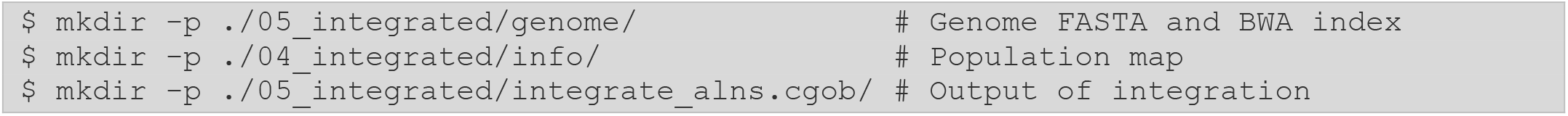

Next, we align the catalog generated in the *de novo* assembly to this *C. gobio* reference genome (code not shown) and then we can integrate the alignment positions into the *de novo* catalog, storing the results in `integrate_alns.cgob`:

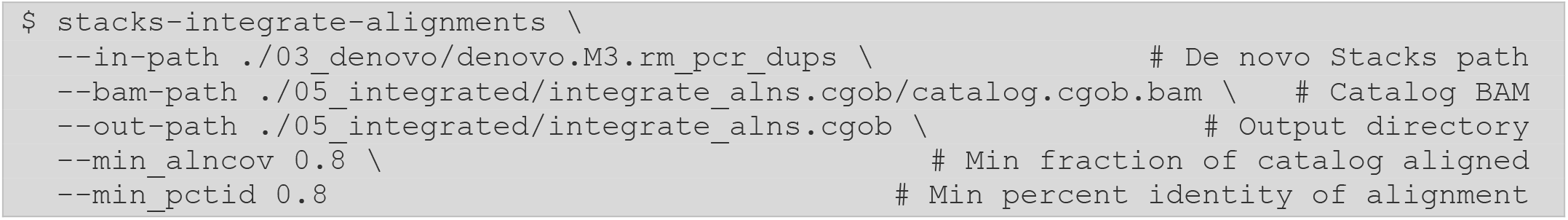

The two filters we employ, minimum alignment coverage (`--min_alncov`) and percent identity (`--min_pctid`), will make sure we avoid spurious alignments of *de novo* loci.

Once `stacks-integrate-alignments` is complete, we then execute the `populations` program on the new catalog in `integrate_alns.cgob` and we are free to proceed with standard filtering, calculating summary and divergence statistics, and exporting data for further analysis.

The strength of the *integrated* results will be related to the evolutionary distance and quality of the heterospecific reference used. Here, with *C. gobio*, many of our loci do not align; starting with 167,698 *de novo* loci, the majority did not align or were filtered, yielding ∼8k loci for analysis. Of course, for demonstration purposes, we used very strict filters and a somewhat distant genome. If multiple high-quality genomes are available, a good approach would be to align the *de novo* catalog to several of them to determine which of the heterospecific assemblies retains the highest proportion of aligned loci.

### 2.8. Populations

In the final analysis stage, the `populations` program puts the data into a population genetics context; in the prior analysis stages, the Stacks pipeline has considered all individuals as a single metapopulation, but now Stacks will divide samples into groups, which may represent geographical populations, divisions according to phenotype, sex, or generation (e.g., F0 and F2) in the case of a genetic map, or species in the case of a phylogenetic analysis. The `populations` program will calculate the frequencies of the observed alleles, according to these divisions, alongside several measures of population diversity and divergence. Importantly, `populations` can apply filters to the data, such as removing alleles and genotypes not observed in the majority of the samples and can export the results for further analysis in many standard file formats (e.g., VCF, GenePop, and STRUCTURE, among others).

The `populations` program is designed to be applied multiple times, independently, over the catalog of loci and genotypes, which were generated by the core pipeline – regardless of if it was generated *de novo* or with a reference genome. In other words, loci are assembled and genotyped once, after which `populations` can be run several times to manipulate, subset, and export the data according to the needs of a given analysis.

We will show an example of a populations command to filter the data generated *de novo* in Section 2.5.2. After the core pipeline execution, the catalog contains all the loci in the metapopulation, including erroneous loci produced by artefacts from the molecular protocol and from noise in the underlying species genome (e.g., repeat elements). A common practice is to only include loci which are present in a fraction of the individuals — with the independent assembly of a locus in multiple individuals serving as evidence that a locus is real. Similarly, to differentiate between real polymorphisms in the population and variation produced by molecular and sequencing error, genotypes can be filtered to only include variant sites present in some fraction of individuals. To begin, we will apply a baseline set of filters to our data to assay how many loci and variant sites are retained and are therefore usable in other downstream analyses. We want to keep loci observed in both populations and present in 50% or more of individuals in each population, and only keep variant sites where the minor allele was observed at least 3 times in the metapopulation (which guarantees they are observed in a minimum of 2 individuals, excluding erroneous SNP calls made only in single individuals). In addition, we want to run this analysis excluding the four samples which we previously characterized as having low quality.

Within the directory of our *de novo* Stacks run we will create a new output directory for this specific `populations` run:

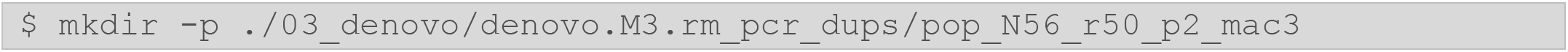

We can then create our `populations` command. First, we specify the path to the base Stacks run we will use for the analysis (`--in-path`). Here, we are using a *de novo* Stacks run, but the analysis can be equivalently performed with reference-aligned or integrated data. Next, we specify the path to our new, filtered population map file (`--popmap`), and the path to our newly created output directory (`--out-path`). We then apply filters, processing only the loci present in at least 50% of samples in a population (`--min-samples-per-pop`), present in both populations (`--min-populations`), and only keeping variant sites with a minimum allele count of three (`--min-mac`). We also determine which loci and variant sites are in Hardy-Weinberg Equilibrium (`--hwe`). Several other options can be found in the Stacks documentation.

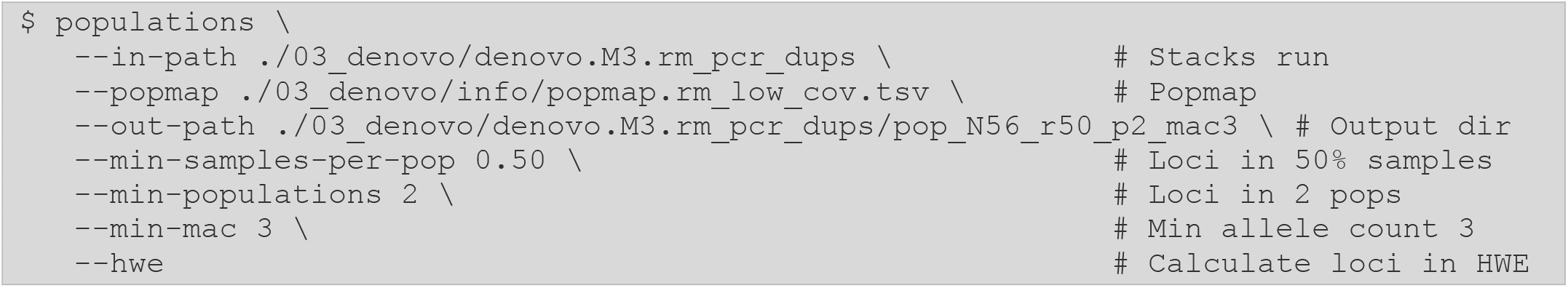

#### 2.8.1. Assessing the populations results

Once the `populations` run is complete, we can evaluate our results by checking the `populations.log` file.

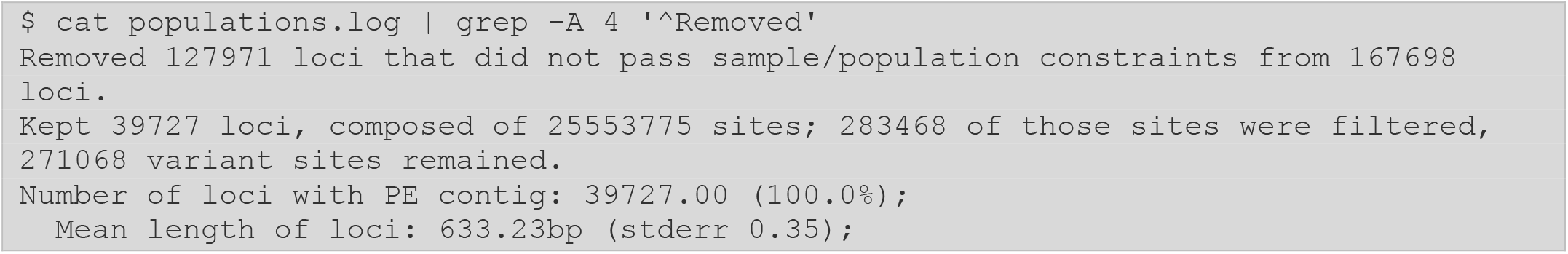

First, we want to check the number of kept loci. We see that, using these filtering parameters, we keep a total of 39,727 loci in this *de novo E. maclovinus* dataset. At first glance, this looks like a small fraction of the 167,698 loci present in the catalog. However, we must remember that the size of the catalog is not representative of the real data as it also contains error loci originating from all samples. Going back to our simulations, we identified around 52k *SbfI* RAD loci in the *E. maclovinus* genome. While this number is higher than the 40k loci seen here, recall that 1) not all loci in the genome will be properly assembled (i.e., those in repeats and recent paralogs), 2) the empirical data contains noise and technical error from the library preparation and sequencing, and 3) that 40k represent the fraction of loci kept after applying moderate filters to the data. The 40k loci kept at this level of filtering stringency should be sufficient for most evolutionary applications, including the 271,068 filtered variant sites (demonstrating that most RAD loci produced more than one SNP). The program also reports to us the mean length of assembled loci (∼630bp). As we increase our filtering stringency, we will typically see this mean length increase.

An additional source of information from the `populations` execution is the `populations.log.distribs` file. This file contains a number of diagnostic distributions, many of them presented both before and after filtering, demonstrating the effects of the chosen filtering parameters. We can view the distribution of the number of samples per locus as well as the number of SNPs per locus. Again, using `stacks-dist-extract` we can pull them out one at a time, first the number of samples per locus, before filtering:

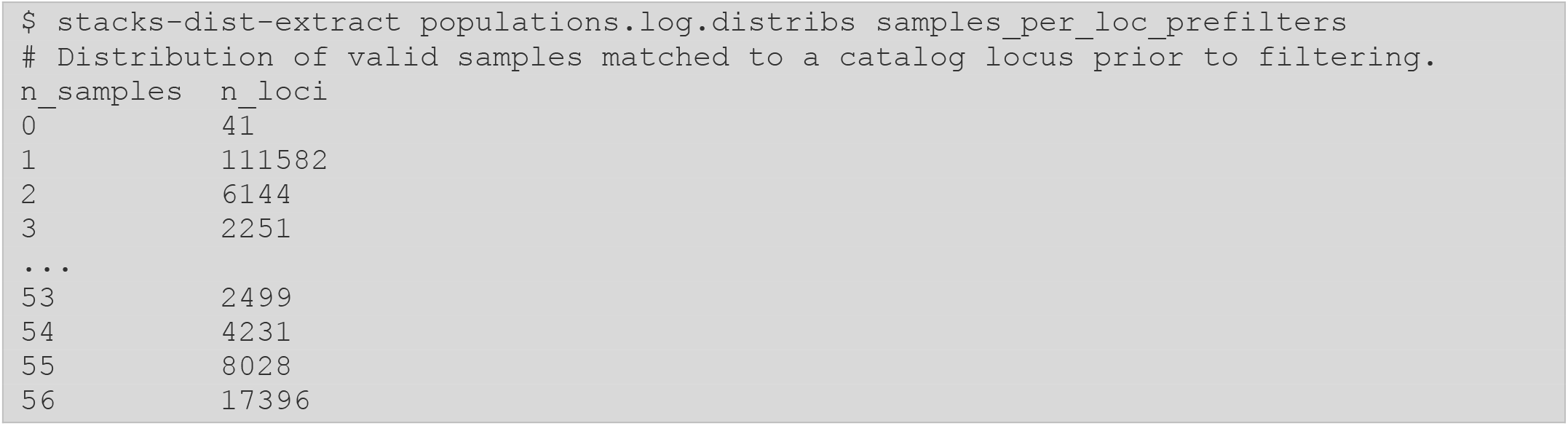

And we can view the number of SNPs per locus, again before filtering:

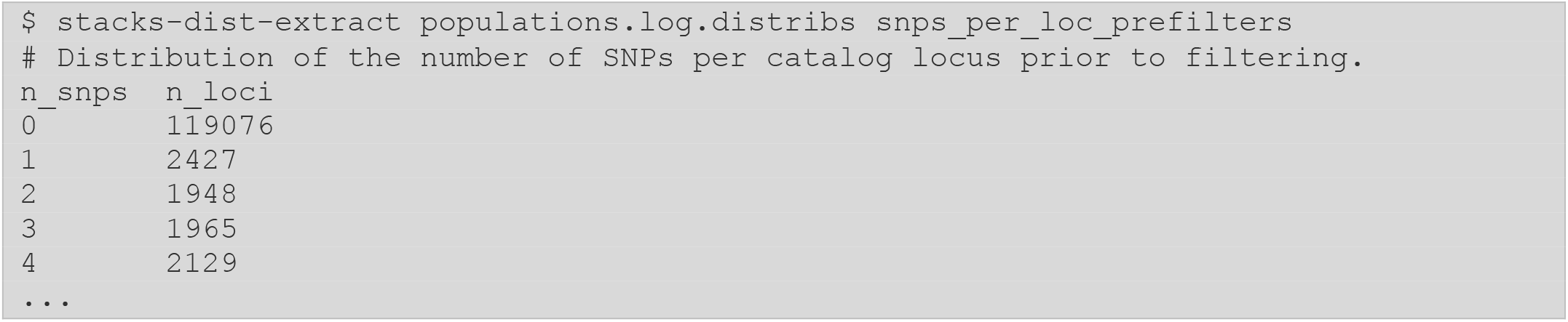

After filtering, it is clear that we have removed loci that don’t meet our 50% threshold, as there are no entries for less than 29/56 total samples (51% of our samples):

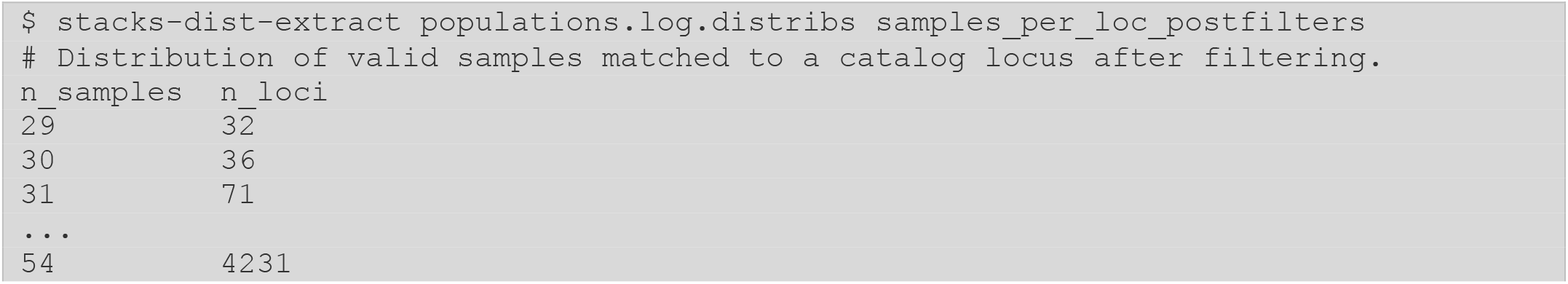
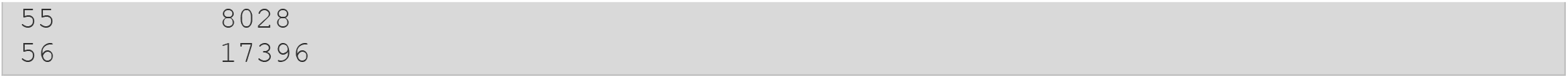

Finally, we can evaluate the amount of data per sample, including the number of missing loci per sample, or the number of missing variant sites per sample. Here, as an example is the number loci per sample:

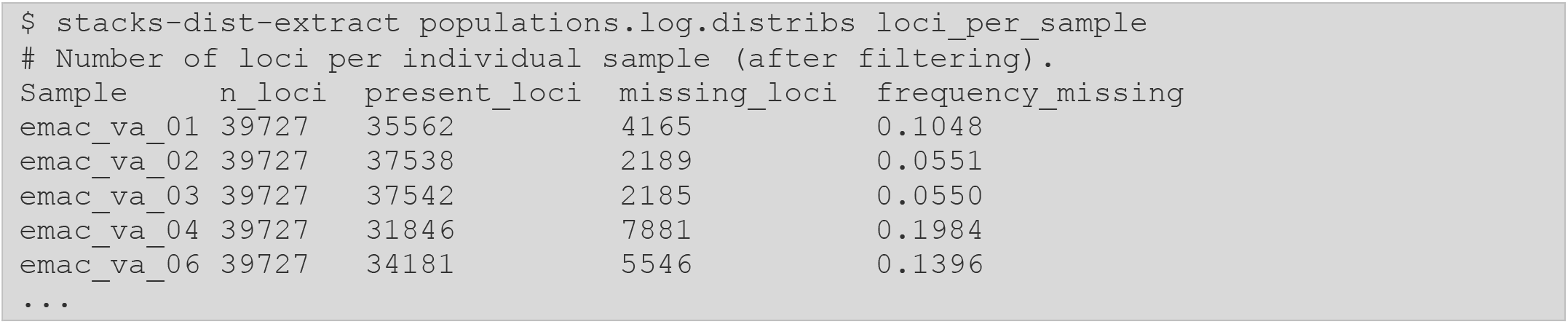

These outputs allow us to see how different individuals in the analysis are performing (e.g., missing data per locus or SNP) as well as evaluating the data set as a whole (e.g., samples per locus) and the underlying biology of our system (SNPs per locus). Two more `populations` outputs provide additional, biologically related information for the system under study. The `populations.sumstats_summary.tsv` file contains a summary of the number of variant sites, heterozygosity, and related measures per population, while the `populations.sumstats.tsv` file contains the same detailed measures per SNP per population.

It is important to note that the filtering parameters shown here are only just a base example of the functionality of the `populations` program. At this point, the onus of interpretation moves from the effects of the molecular protocol and sequencing run to the underlying biology. It is up to the researcher to determine what level of SNPs or loci are appropriate and these values will differ widely, for example searching for population structure in closely related populations versus building phylogenetic trees among distantly related taxa will present very different requirements. Researchers should not forget that the filtering parameters are not the goal of the analysis. Instead, practitioners should shift their thinking to ask: What is required to answer the biological questions proposed in the experiment?

In the following sections, we provide several possible routes for further analysis, beyond Stacks itself.

#### 2.8.2. Post-analysis directions after Stacks

Given a set of RAD loci, and their endemic SNPs, nearly any population genetic analysis is possible. However, different assumptions apply to the underlying data in different analyses, and we can use the populations program to prepare different sets of loci and or SNPs for these analyses. We provide several examples below.

*Note! In the examples below we apply specific filtering parameters for each of our analyses related to the quality and coverage of the E. maclovinus dataset and based on the requirements of the downstream application. Specific filters should be tested and verified to be appropriate for each independent dataset*.

##### SNP-based population structure

One of the most common population genetic analyses is to search for population structure, or to understand how many genetic groups are present in a set of individuals. This can be done via a PCA analysis, or by running the program STRUCTURE (Pritchard et al. 2000) or one of the newer derivatives. To determine population structure, we only need a small number of SNPs, and we don’t need genomic coordinates for those SNPs (they can be from a *de novo* analysis). However, since we want to understand the underlying neutral relationships between individuals, we want to avoid outlier SNPs whose allele frequencies may be driven by selection – choosing SNPs in Hardy-Weinberg Equilibrium (HWE) can ameliorate this concern. Further, multiple SNPs from the same locus are by definition linked, so we only want to choose one representative SNP from each locus. The `populations` program will calculate how far out of HWE a locus is, providing a p-value, and this data is included for individual SNPs (`populations.sumstats.tsv`) and for loci (`populations.hapstats.tsv`). In addition, we can instruct `populations` to only include a single SNP per locus (chosen after filtering is complete) and we can supply a *whitelist* of loci – a specific set of loci to include, enabling us to randomly select, for example, 5,000 loci for the STRUCTURE analysis.

First, we will create a whitelist file, based on the loci recovered in the previous `pop_N56_r50_p2_mac3` run. We will randomly select 5,000 loci that are in HWE for both populations. Here we employ a shell pipe to combine Stacks output data to generate our list of loci:

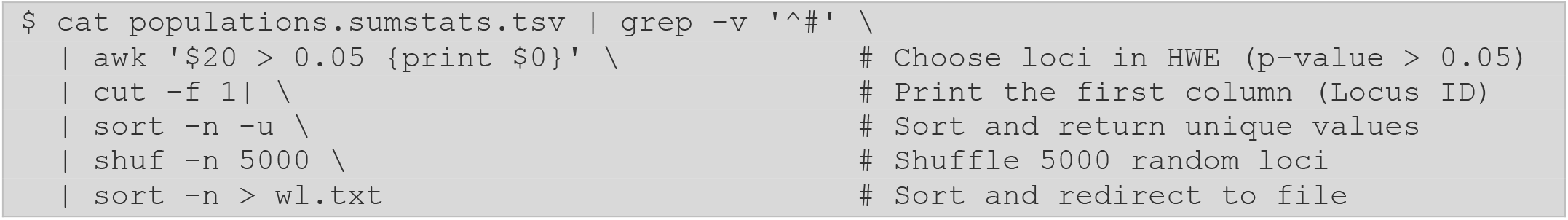

We will also create a new directory to output this new run:

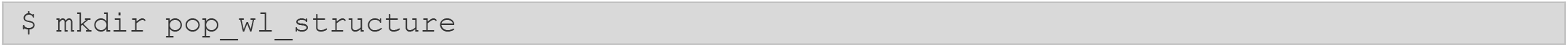

Once we have created our whitelist, we can form the populations command. Note we no longer need filters since the list of loci in our whitelist originated from an already-filtered set of loci.

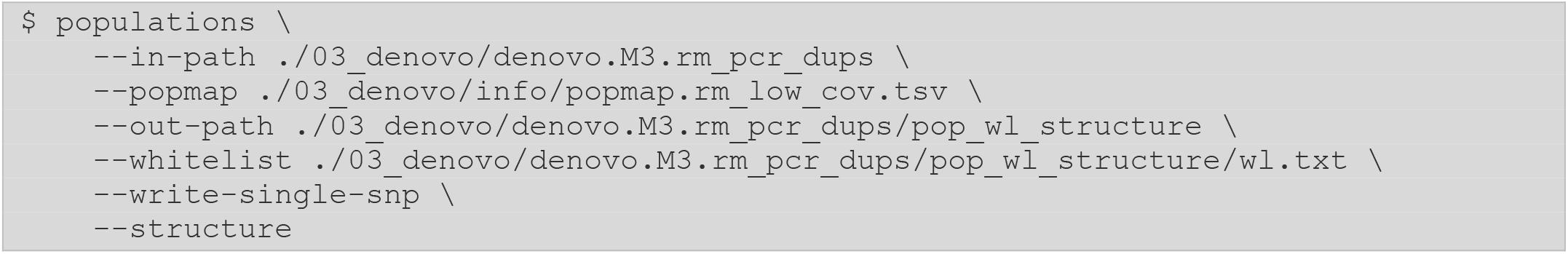

After the populations run is complete, we will find a `populations.structure` file in the output directory, which can then be loaded into STRUCTURE (after removing the Stacks timestamp comment on the first line and modifying the population identifiers). We can visualize the output for our *E. maclovinus* dataset for K=3 (Fig. 5A), where we observe a substructuring of samples within the Valdivia population (`va`). Little to no substructuring is observed within the two geographic sites.

**Figure 5.**
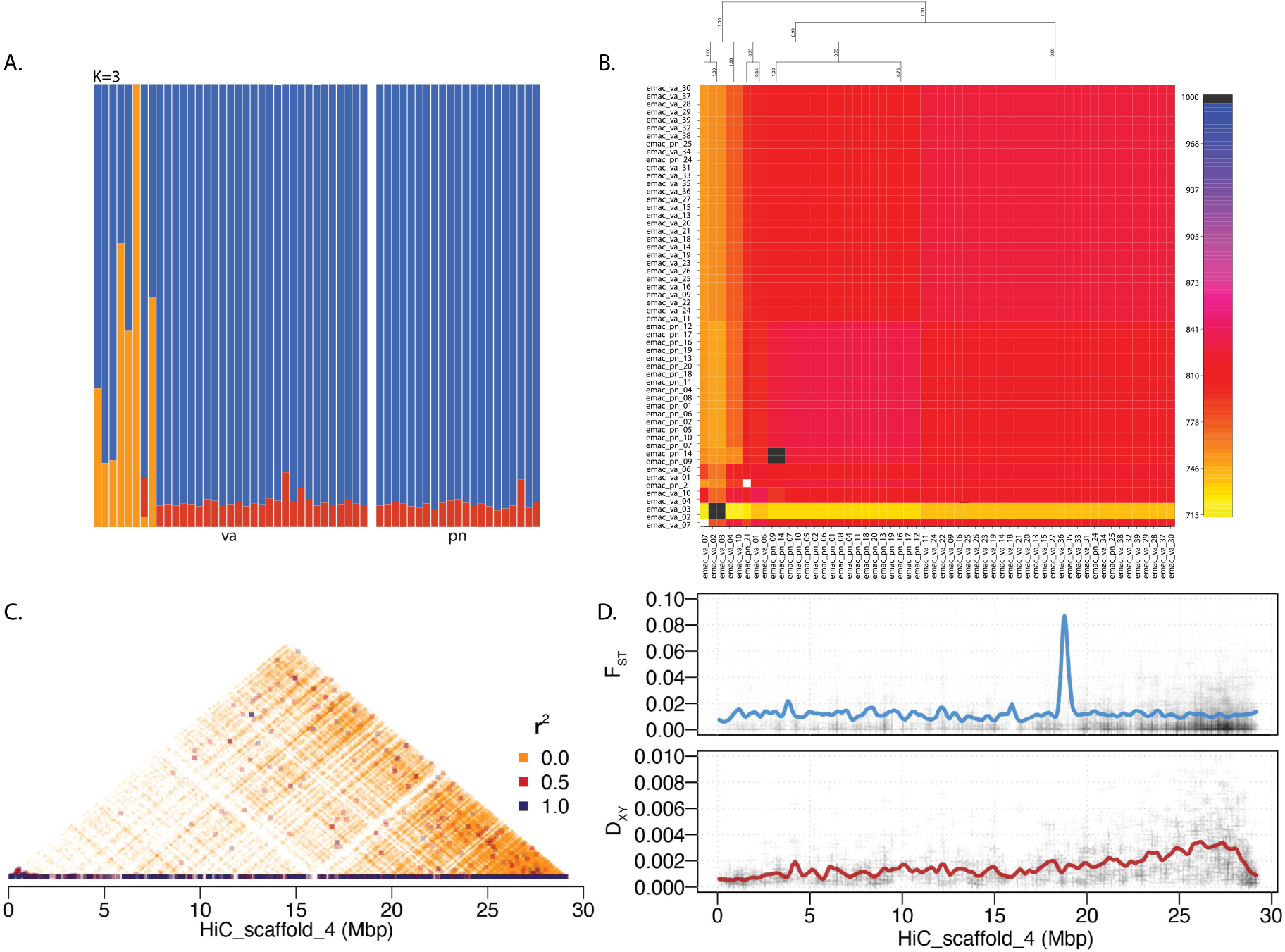
Examples of downstream applications of the populations output. (A) SNP-based assessment of population structure using STRUCTURE, with a K=3, for the Valdivia (`va`) and Puerto Natales (`pn`) *E. maclovinus* populations. (B) Haplotype-based assessment of population structure using RADpainter/fineRADstructure. Heatmap shows population clustering based on sample co-ancestry. (C) Pairwise linkage disequilibrium (measured by the correlation coefficient, *r*^*2*^) in the *E. maclovinus* HiC_scaffold_4. This was calculated by phasing SNPs obtained via a reference-based analysis. (D) Genetics divergence along the *E. maclovinus* HiC_scaffold_4. Top plot shows relative divergence (F_ST_). Black crosses show per-site F_ST_values, while the blue line shows a smoothed average. Bottom plot shows absolute divergence (D_XY_), with the black crosses showing per-locus D_XY_values and red line showing a smoothed average.

##### Haplotype-based population structure

An alternative method for assessing population structure uses haplotypes instead of the single SNPs used by STRUCTURE. A haplotype approach maximizes the biological information encoded by each marker—single SNPs can only describe two alleles present in the metapopulation, while haplotypes can describe several alleles. This increase in information content by markers often allows for an increase in resolution both when identifying differentiation between populations and when assigning individuals to a population (Leitwein et al. 2020; Bootsma et al. 2021).

Identifying population structure using RADseq-derived haplotypes can be performed using `RADpainter`and `fineRADstructure`(Malinsky et al. 2018), a pair of sister programs designed for the generation of a co-ancestry matrix derived from RADseq haplotypes, followed by the inference of populations among a group of individuals based on their co-ancestry. This analysis was specifically design around RADseq data, thus can take full advantage of the haplotypes natively produced by Stacks.

Similar to STRUCTURE, Stacks can export data directly in the `RADpainter`/`fineRADstructure` format (`--radpainter`). Since the software is designed to work around a larger number of markers, it is not necessary to subsample loci using a whitelist. Instead, we can just apply our filters directly (`--min-samples-per-pop 0.75`, `--min-populations 2`). In addition, we tell populations to filter data by haplotypes (`--filter-haplotype-wise`), which applies the missing data filters to the whole locus, instead of applying them to the individual variant sites. This approach will prune the variant sites with the highest proportion of missing data in the locus, in turn guaranteeing the least possible missing data within the kept haplotypes. This option is recommended for all applications based on the haplotype outputs from `populations`.

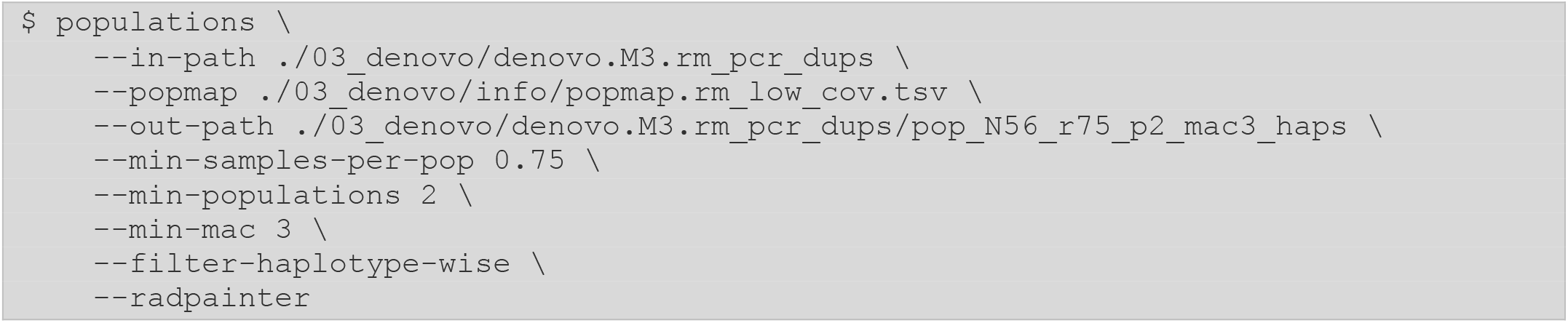

The generated `populations.haps.radpainter` file can be imported directly into RADpainter/fineRADstructure for analysis. Applying this method to our *E. maclovinus* dataset (Fig. 5B) we observe that there is a fraction of `va` samples clustering separately from all other individuals (see Section 2.9.3). In contrast to STRUCTURE, the extra information present in the haplotypes does allows for the sub-clustering of `va` and `pn`individuals.

##### Divergence statistics

An alternative analysis route involves looking for divergent loci between populations. This can include traditional measures of F_ST_and is again calculated at the SNP level. It is possible to calculate F_ST_without a reference genome, typically plotting loci with high levels of heterozygosity against those loci with high values of F_ST_(Allendorf et al. 2010). However, once these loci have been identified, there are not a lot of further analytical options. Given a reference genome, however, `populations` can identify SNPs that are co-linear on particular chromosomes indicating regions of divergence between populations. Below, we run `populations` on our reference-aligned dataset, and we employ several filters to export loci present in 80% of samples (`--min-samples-per-pop 0.80`) and present in two populations (`--min-populations 2`), and removing variant sites with a minimum allele frequency below 5% (`--min-maf 0.05`). We then ask for F statistics to be calculated (`--fstats`) and for the software to *smooth* the values of F_ST_, giving us a moving average of divergence along each chromosome:

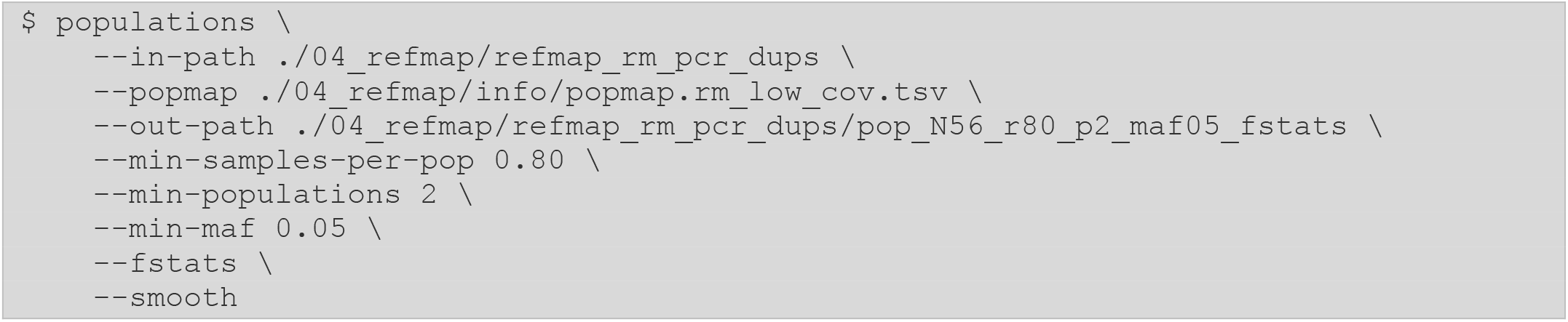

Once `populations`is complete, an output file will be created for each pair of populations in the analysis, containing per-SNP divergence statistics. For the *E. maclovinus* dataset, this output file is named `populations.fst_va-pn.tsv`for the `va`and `pn` populations. In the case of genetic divergence, there are also several statistics that are applied at the locus/haplotype level, including absolute divergence, D_XY_, and locus-level forms of F_ST_including Φ_ST_and F_ST_’. These multiallelic statistics can provide a stronger signal, since the alleles can be distributed across multiple populations in more configurations than biallelic SNPs. D_XY_also provides a more robust divergence statistic at more distant time scales as it is directly proportional to the inter-population coalescent time without being biased by allele frequency (Cruickshank and Hahn 2014). And may be better for the identification of islands of divergence, as well as providing evidence for additional models for divergence with ancient or non-existent gene flow (Irwin et al. 2018). These files are output by `populations` in a similar way, including smoothed values, in `populations.phistats_va-pn.tsv`.

The output of these divergence analysis can then be used for several applications, one example is plotting genetic divergence along specific chromosomes. Here we can see the relative (F_ST_) and absolute divergence (D_XY_) between the `va` and `pn` *E. maclovinus* populations over the chromosome-level scaffold, `HiC_chromosome_4`(Fig. 5D).

##### Linkage disequilibrium

When reference-aligned RADseq data are available, it can additionally be used to generate measures of linkage disequilibrium (LD) along the chromosomes to identify patterns of outlier recombination and linkage. Before we can perform any LD calculations, we first must statistically phase our variant sites into chromosome-level haplotypes. Since the phasing process imputes the genotypes for any missing sites, we will use very stringent filtering parameters (loci present in over 90% of individuals) to guarantee we export the least number of missing genotypes. Additionally, we will apply the `--ordered-export` flag to `populations` to ensure sites are exported sorted by their genome coordinates, and in the case of overlapping RAD loci, only one representative is exported, guaranteeing compatibility with external software.

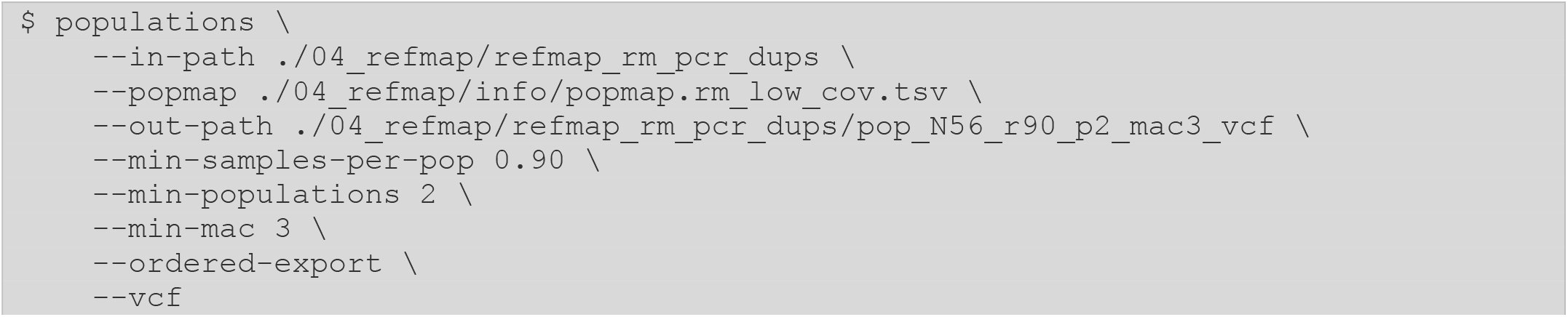

The exported VCF can then be phased using BEAGLE (Browning and Browning 2007; Browning et al. 2018), after which LD calculations can be performed using VCFTOOLS (Danecek et al. 2011). Once phasing is complete, measures of LD can be plotted along the chromosomes to identify regions displaying extended linkage, as shown in (Fig. 5C). Additionally, other LD-derived measures, such as extended haplotype homozygosity, can be computed on the chromosome-wide, phased results using, for example, the REHH (Gautier et al. 2017) R package.

### 2.9. Triaging negative results

When poor results occur with a RADseq experiment, often the practitioner may try to *rescue* the data by applying *strict* analytical practices (this is inherently very subjective, hence the italics). Based on our experience in RADseq analysis, here we consider three common questions that result from a poor analysis.

### 2.9.1. Should I trim data?

A common procedure of many analysis pipelines for the processing of short-read Illumina data is the trimming of sequencing reads to remove fragments of the read with low quality and/or containing adapter sequences. While this produces datasets of uneven read length, the downstream effects of coverage are ameliorated by the random position of the shotgun reads along the genome (i.e., the trimmed portion of the reads will not always originate from the same position on the genome). This is, however, not the case for RADseq data – all reads from a single locus are anchored to the cutsite at the 5’-end of the locus. Trimming RADseq reads will therefore result in a localized decrease in coverage at the 3’ of the locus, which can have downstream effects on genotyping (since the calls are dependent on available coverage) and the assembly of contiguous loci (since the single- and paired-end contigs are merged based on their overlapping sequences). Counterintuitively, trimming low quality data may produce even worse results.

While not advised for the reasons described above, the Stacks pipeline can analyze trimmed data, as the gapped aligner in `ustacks` will handle uneven reads as if containing gaps. However, the software might throw out very short reads completely if they are unable to be matched to a putative locus. Sequencing a dataset to low coverage can exacerbate these behaviors, since the removal of reads results in a proportionally larger reduction of coverage across the whole locus, not just the 3’ end (Fig. 6A). The overall reduction in coverage subsequently results in a further increase of missing genotypes locus-wide (Fig. 6B).

**Figure 6.**
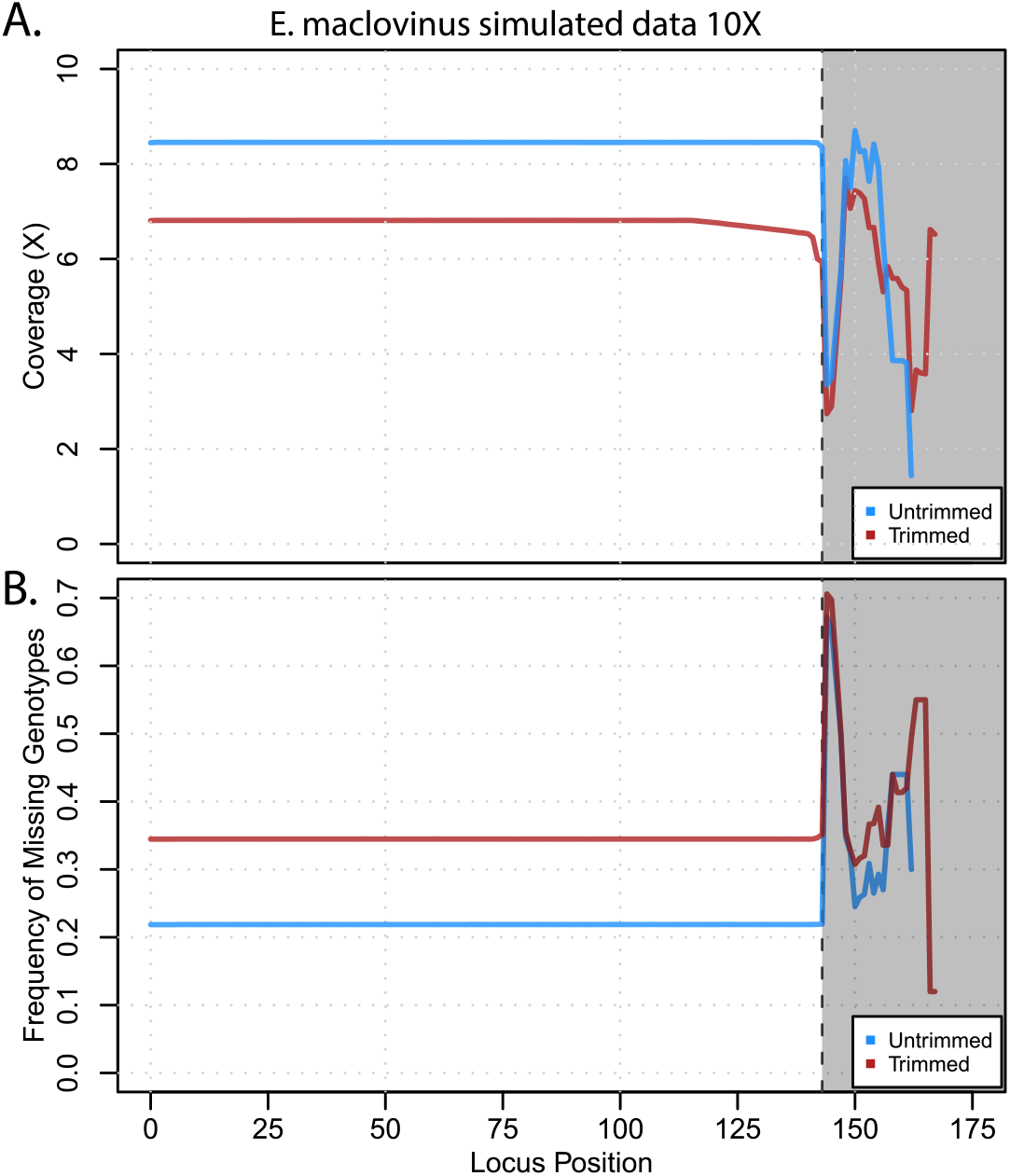
Effects of read trimming on *de novo* locus assembly at low coverages. The results are shown for a simulated RADseq dataset, calculating values by averaging across all assembly loci. (A) Change in coverage along the single-end (forward read) region of the RAD locus. Grey dashed line shows the forward read length after barcode removal (143bp). Grey box shows values for regions beyond the read length produced by indels between samples. The untrimmed dataset (blue) shows constant coverage across the locus, while the trimmed dataset (red) shows a decrease in coverage along the 3’ end of the locus. (B) Frequency of missing genotypes along the single-end region of the RAD locus. Grey dashed line shows the forward read length (143bp). Grey box shows values for regions beyond the read length. The untrimmed dataset has a constant proportion of missing genotypes of around 20% along the locus, resulting from the low coverage. The trimmed dataset, while also having a constant value along the locus, shows an increase in missing genotypes, around 35%.

Instead of trimming the datasets, the approach taken by Stacks during `process_radtags` is focused on the processing and discarding of whole reads instead of just partial fragments. As seen in Section 2.4, with proper initial sequencing coverage, whole reads can be filtered from the dataset without negative impacts on downstream coverage and genotypes. Reads with poor sequencing quality are removed altogether with the `--quality` flag and checking for and discarding reads with adaptor fragment sequences can be done by using the `--adapter_1` and `--adapter_2` options. If coverage is low and adaptor is present, a better strategy may be to trim all reads to an even length, with `-t`. However, these options should only be applied after proper assessment of the quality of the input data to prevent the discarding of excess sequence.

#### 2.9.2. Controlling the stringency for calling genotypes at low coverage

RAD practitioners often want to apply a hard depth of coverage limit to assembled loci. Stacks does not provide this type of filter primarily because it results in arbitrary cut offs – often removing loci that are statistically supported by their genotype calls, and/or not removing loci whose depth is inflated by PCR duplicate reads. Instead, if a user wants to have higher confidence in genotyping results (for which depth of coverage is used as a proxy) they should increase the level of evidence required for the genotyping model to make a call. Reducing the alpha value to the genotyping model (e.g., from 0.05 to 0.01) requires more evidence, e.g., reads or depth of coverage, for a call to be made.

Returning to our original results from the *de novo* analysis (Section 2.5), after genotyping was performed in `gstacks` with the default parameters, and moderate filters were applied by the `populations` program (Section 2.8), the analysis produced:

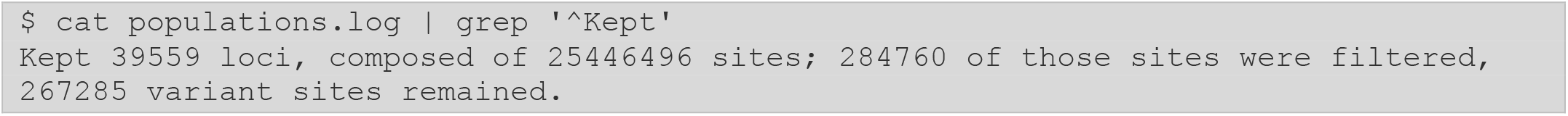

We had 606,487 variant sites on this dataset by genotyping with default parameters. We then reran `gstacks` with `--gt-alpha 0.01`, to be more stringent in our genotype calls:

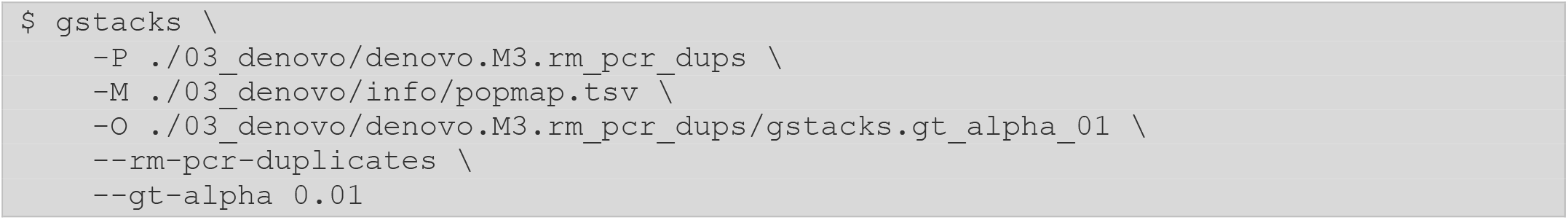

We then rerun `populations`, applying the same filters as before (loci present in 50% of samples and two populations, variant sites observed three times in the catalog):

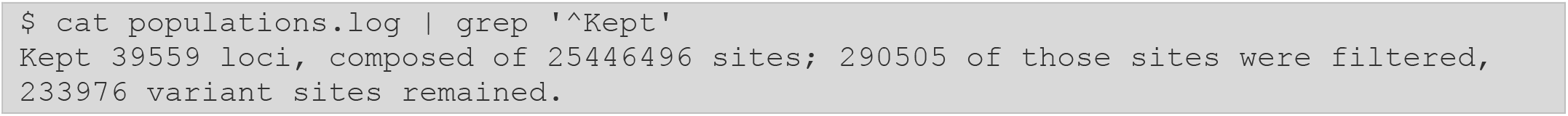

We retain the same number of loci (40k) and the same number of total sites (25M), but the original run retained 267k variant sites, while our new run with more stringent genotyping retains only 234k. This shows that we likely have 33k variant sites with low coverage which could not be genotyped after making `--gt-alpha` more stringent. However, this does indicate that the variant sites that are retained after this increase in genotype stringency are called very high confidence.

These changes in genotyping stringency are a good approach for dealing with the effects of low coverage. Instead of applying an arbitrary coverage cutoff, users should simply let the genotyping algorithm asses the available evidence. The result will likely be a tradeoff between the magnitude and confidence of genotypes, with the final decision on stringency cutoff being dictated by the user according to the requirements of the experiment.

#### 2.9.3. Removing Bad Apples

Cerca et al. (2021) described a method for the optimization on RADseq data based on the removal of samples with high missing data, or *bad apples* (Cerca et al. 2021). Briefly, a Stacks catalog is built (either *de novo* after parameter optimization or using a reference genome) after which the researcher identifies samples containing a high proportion of missing genotypes (described in Section 2.8.1). These samples drive a significant proportion of the catalog-wide fraction of missing data, and their removal improves overall retention of loci and genotypes after filtering, as well as minimizing biases derived from missing genotypes.

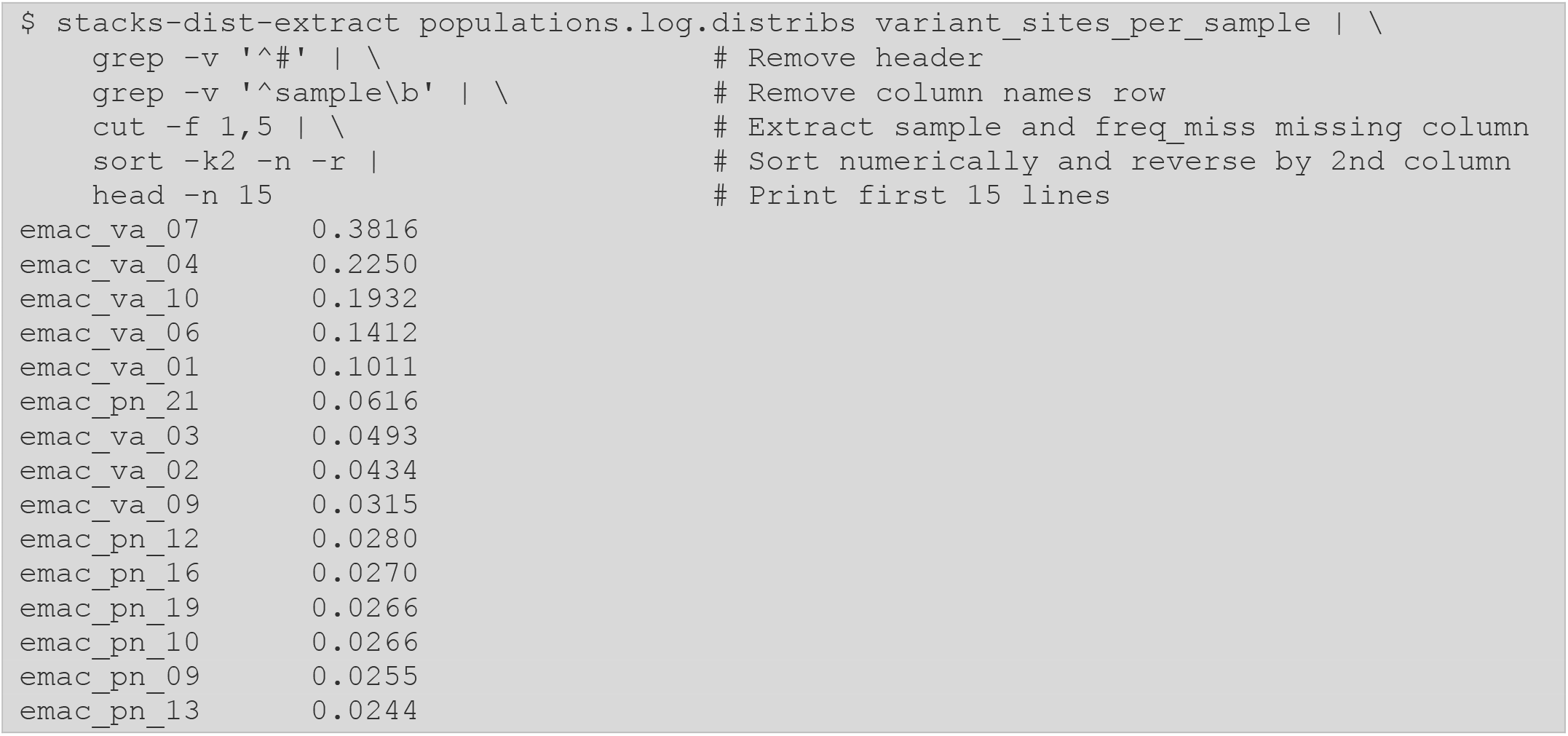

When viewing our *E. maclovinus* dataset in the light of bad apples, we do identify a fraction of individuals containing a higher proportion of missing data than the other samples in the analysis, even after the removal of some samples following raw read processing (Section 2.4.1) and catalog construction (Section 2.5.3). For example, when reviewing the `RADpainter`/`fineRADstructure` analysis performed previously in (Section 2.8.2), we notice that the proportion of `va` samples clustering separately from all other individuals are also the ones with the elevated proportion of genotypes:

While the median proportion of missing data is less than 2% (median: 0.01845, mean: 0.03774), notice that some samples show outlier fraction of missing genotypes. This subset of samples can be classified as bad apples and removing them from subsequent analysis (by removing them from the population map file) can improve downstream analysis. For example, removing the apparent missing data bias in the results from `RADpainter`.

## 3. Appendix

Working example scripts referenced in this protocol can be found online at BitBucket in a Git repository: https://bitbucket.org/CatchenLab/mimb-stacks2-protocol. We list the available scripts below.

### 3.1. Plot library demultiplexing and processing

The file `01_process_radtags/process_radtags_stats.R` uses the log reported by `process_radtags` and calculates several summary statistics on the proportion of samples in the library, number of total reads, and percentage of total reads kept per-sample. It also generates several plots showing the generated distributions. The input file can be obtained using:

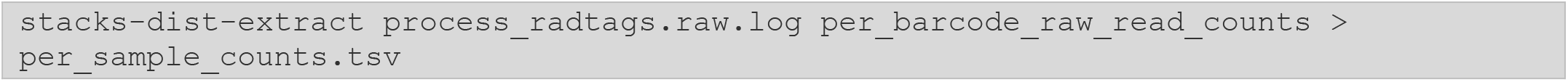

### 3.2. *De novo* parameter optimization shell loop

The file `02_param_opt/param_opt_loop.sh` loops over the values of 1 through 12 and uses the number in each iteration as the corresponding value for `ustacks -M` and `cstacks -n` for the Stacks *de novo* pipeline.

### 3.3. gstacks coverage R script

The file `03_denovo/gstacks_stats.R` uses the log reported by `gstacks` and calculates several summary statistics on sample coverage and PCR duplicates. It also generates several plots showing the generated distributions. The input file can be obtained using:

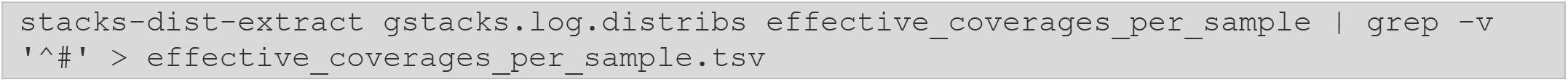

### 3.4. BWA alignment shell loop

The file `04_refmap/bwa_samples_loop.sh` loops over each sample in the provided population map file, and for each sample aligns the paired reads with `bwa mem` and processed alignments using `samtools view/sort`. For each iteration of the loop, the sample name present in the popmap is used as the prefix from which the path the input reads and output BAM files are constructed.

